# VAETracer: Mutation-Guided Lineage Reconstruction and Generational State Inference from scRNA-seq

**DOI:** 10.64898/2026.01.19.700238

**Authors:** Linnuo Pan, Kaiyu Wang, Luonan Chen

**Author notes:** Co-first Author. Corresponding Author: Luonan Chen.

## Abstract

Somatic mutations accumulate with cell division and are key to understanding tumor evolution. While single-cell RNA sequencing (scRNA-seq) can effectively capture somatic mutations in the 3′ untranslated region (3′ UTR), enabling its use for lineage tracing, these data are inherently noisy and exhibit high false-positive rates. To address this limitation, we propose VAETracer, a deep learning framework that is able to reconstruct cellular lineages by extracting cellular generation index (CGI) from mutation profiles of 3′ UTR in scRNA-seq data, thus enabling the inference of developmental trajectories without relying on noisy mutation signals (https://github.com/Kaiyu-W/VAETracer). There are two core components for VAETracer: (1) scMut module, which infers the CGI directly from sparse and noisy mutation matrices using our cumulative mutation model (CMM), effectively bypassing error-prone phylogenetic tree topologies caused by mutation noise, and (2) MutTracer module, which predicts ancestral and future latent cellular states or gene expressions at the single-cell level by integrating CGI information with measured transcriptomes. Validation on simulated and real tumor datasets shows that this method can accurately reconstruct clonal relationships, quantify tumor progression, and effectively infer unmeasured cellular states using only the measured scRNA-seq data. This work provides a new computational tool for extracting lineage information and revealing tumor evolution directly from widely available RNA-seq data.

## Backgrounds

Characterization of somatic mutations at single-cell resolution is essential for understanding tumor evolution, as mutations accumulate progressively across cell generations, driving genetic heterogeneity, cellular plasticity, and clonal diversification during cancer progression(McGranahan and Swanton, 2017; Neftel et al., 2019; Nowell, 1976). During tumor progression, somatic mutations are progressively acquired across successive cell generations, providing a natural temporal axis along which clonal evolution and transcriptional state transitions unfold(Black and McGranahan, 2021; Funnell et al., 2022; Vogelstein et al., 2013).

A variety of experimental and sequencing-based approaches have been developed to interrogate tumor evolutionary dynamics, as lineage tracing in the sequencing era has become a central challenge in developmental and cancer biology(Shapiro et al., 2013; Wagner and Klein, 2020). Single-cell genome sequencing (scDNA-seq) has been most widely used for lineage tracing based on copy number variations (CNVs)(Andor et al., 2020). scDNA-seq also offers somatic mutations at single-cell resolution (Pellegrino et al., 2018; Weber et al., 2023), but its practical application is constrained by limited scalability, allelic dropout, and amplification-induced technical noise(Petti et al., 2019; Valecha and Posada, 2022). Similarly, mitochondrial DNA (mtDNA) variants have been investigated as endogenous lineage markers(Ludwig et al., 2019; Xu et al., 2019), while their stochastic segregation during cell division and lack of information on nuclear genomic alterations limit their utility and reliability for reconstructing clonal relationships(Li et al., 2025b).CRISPR/Cas9-based lineage tracing has enabled high-resolution reconstruction of lineage relationship (McKenna et al., 2016; Raj et al., 2018; Simeonov et al., 2021). However, these approaches rely on engineered barcodes rather than endogenous mutations and primarily encode relative lineage relationships, with their application further limited by experimental complexity, cellular toxicity, and conditional requirements(Salvador-Martinez et al., 2019; Weinreb et al., 2018). Despite these advances, jointly resolving lineage relationships, temporal progression, and transcriptional state changes at single-cell resolution remains challenging.

In addition to experimental lineage-tracing methods, single-cell RNA sequencing (scRNA-seq) has been used to infer lineage through copy number variation (CNV) profiles derived from transcriptomic data(Li et al., 2022; Wei et al., 2024). Beyond CNV-based approaches, high-throughput scRNA-seq also enables the detection of expressed somatic mutations, particularly in the 3′ untranslated region (3′ UTR), thereby offering a dual readout of genetic and transcriptional states at single-cell resolution(Anderson et al., 2022; Bai et al., 2025; Chen et al., 2016; Vu et al., 2019).

However, mutation detection from scRNA-seq is challenged by sparse coverage, allelic drop-out, RNA editing, and sequencing artefacts (Kharchenko et al., 2014), resulting in mutation matrices that are highly sparse, noisy and prone to false positives, particularly in 10× 3′ scRNA-seq datasets. While several methods have been developed to identify high-confidence somatic mutations from scRNA-seq data, such as SComatic(Muyas et al., 2024) and RESA(Zhang et al., 2023), focusing on mutations associated with specific cell types or biologically relevant programs. However, the limited yield of such high-confidence calls renders them insufficient for robust downstream clonal-evolution inference. To address this limitation, methods specifically designed to infer clonal structure from scRNA-seq mutations, such as scClone(Bai et al., 2025) and PhylinSic(Liu et al., 2023). Still, these methods typically rely on imputing missing information from neighboring cells, which may introduce biases and reduce resolution in the inference of fine-grained clonal architectures.

Building on these advances and also addressing the limitation of noisy 3′ UTR mutation signals in widely available scRNA-seq data, we proposed a cumulative mutation model (CMM) to infer cellular generation index (CGI) from sparse and noisy mutation matrices. Rather than relying on explicit phylogenetic tree reconstruction, VAETracer uses CGI to provide a generation-aware temporal reference and jointly models mutational and transcriptional features to infer ancestral and future cellular states. By integrating lineage information with transcriptional dynamics, VAETracer enables time-resolved analysis of tumor evolution using widely available scRNA-seq data. Validation on simulated and real tumor datasets demonstrates that this approach reconstructs clonal relationships, quantifies tumor progression, and predicts unmeasured cellular states, providing a unified framework for studying the interplay between somatic mutations and transcriptional plasticity in cancer.

## Results

### Overview of VAETracer

We developed VAETracer (Fig.1), a deep learning–based framework for mutation-informed lineage analysis from scRNA-seq data. The model integrates somatic mutation signals with transcriptional information to support analyses of lineage structure, cellular generation, and gene expression dynamics across tumor evolution.

**Figure 1.**
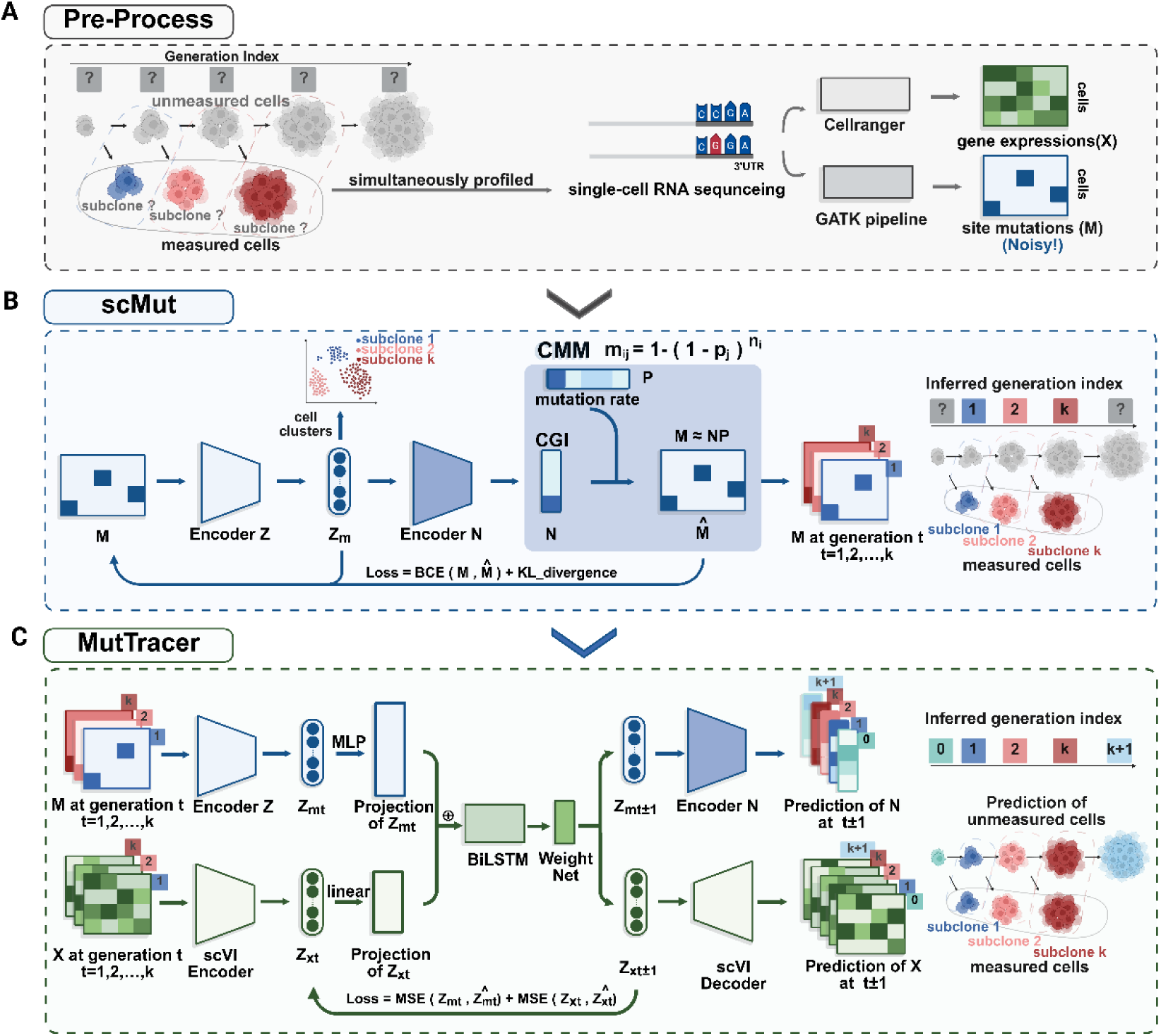
Overview of VAETracer. Workflow of the VAETracer framework consisting of three modules. (A) Pre-processing of scRNA-seq data. Cells from multiple subclones, each corresponding to a specific cellular generation, are simultaneously profiled in a single scRNA-seq experiment and processed through standard analysis pipelines. Gene expression counts (X) are quantified using Cellranger, while site-specific mutations (M) are detected via a GATK pipeline, however the resulting M is inherently noisy due to technical limitations and sparse coverage. (B) scMut: CGI inference under a CMM. The mutation matrix M is first embedded via an encoder to produce a latent mutational representation Z_m_, which can be used to perform clustering of subclones. Another encoder infers the CGI (N), which is used in combination with the mutation rates P by a CMM to estimate the expected mutation probabilities (*M̂*) at each generation. The model is trained using a combination of binary cross-entropy (BCE) loss for mutation reconstruction and KL divergence for latent regularization. Predicted mutations profiles *M̂* are then separated to each generation index t=1, 2, …, *k* according to N. (C) MutTracer: mutation-guided temporal state inference. For each generation index t, the mutational latent representation Z_mt_ is obtained from the scMut encoder Z and further transformed via an MLP projection, while the transcriptional latent representation Z_xt_ is extracted from the gene expression matrix X_t_ using a scVI encoder and projected via a linear layer. These projected representations are concatenated and input to a BiLSTM to capture forward and backward temporal dependencies across generations. A weight network generates dynamic modality-specific weights to separate fused features into mutational and transcriptional components, then predicts Z_mt±1_ and Z_xt±1_, from which the corresponding N _t±1_ and X _t±1_ are inferred via the scMut encoder N and the scVI decoder.

scRNA-seq data are first processed to obtain a unified representation of both gene expression and somatic mutation profiles. In principle, cells directly related along a lineage trajectory provide an ideal system for studying developmental dynamics; however, in practice, harvested cells cannot generate progeny, making it impossible to obtain true lineage relationships experimentally. Consequently, lineage structure must be inferred indirectly, with cellular generation serving as a key proxy for reconstructing developmental progression. Within a single scRNA-seq experiment, cells may originate from multiple subclones corresponding to cellular generations, captured simultaneously at the time of measurement. As scRNA-seq provides only a transcriptional snapshot, generational information is not explicitly encoded and cannot be directly observed. Moreover, not all ancestral or descendant cells are sampled, resulting in a partial and temporally collapsed view of the full lineage. From the measured scRNA-seq reads, we derive a gene expression matrix X and a mutation matrix M, where the mutation signals are inherently sparse due to technical limitations and low allelic coverage. While X captures transcriptional states across cells, M provides complementary lineage-related information that can be used for downstream modeling or phylogenetic reconstruction.

We then developed the single-cell Mutation Parser (scMut) tool, which takes the sparse mutation matrix M as input and is based on a cumulative mutation model (CMM). Under this model, mutations are assumed to accumulate progressively across cell generations, such that the occurrence of a mutation at each genomic site follows a generation-dependent Bernoulli process with site-specific probability p. Specifically, for cell i and site j, the expected mutation probability m_ij_ is defined as

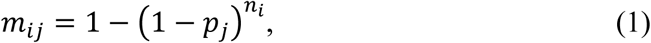

where n_*i*_ denotes the cellular generation index and p_j_ represents the site-specific mutation rate. Actually, as a linear approximation of the right-hand-side for this formulation above (eqn.(1)), it can be expressed by *m*_*ij*_ = *n_i_p_j_*, i.e., a non-negative matrix factorization (NMF) form or M=NP. In other words, as an approximation, NMF can also be applied to M to jointly estimate N and P. Here, we instead adopt a more direct decomposition approach by optimizing the original CMM, which we refer to as generalized NMF (gNMF). In parallel, a variational autoencoder (VAE) is employed to learn a low-dimensional latent representation Z_m_ of mutational profiles, which further constrains the inference of the cellular generation index N. Predicted mutations *M̂* can then be assigned to individual generations index t=1, 2, …, *k* based on the inferred N, allowing downstream analyses to explicitly model generational dynamics. Together, scMut provides a continuous and noise-robust estimate of cellular generational progression directly from sparse scRNA-seq mutation profiles, without relying on explicit phylogenetic reconstruction.

After this, we introduce a unified framework Single-cell Mutation-guided Temporal State Inference (MutTracer) that performs generation-aware joint modeling of mutational and transcriptional dynamics. Cells are ordered into discrete generation index t according to N obtained from the scMut, or by explicitly provided multi-timepoint annotations when available. The aligned expression matrix X is embedded into a low-dimensional representation Z_x_ using scVI(Lopez et al., 2018), while the mutational latent representation Z_m_ is obtained from the scMut. The mutational and transcriptional latent representations are first processed separately through modality-specific projection layers, reflecting their distinct contributions. Specifically, Z_m_ is transformed via a multi-layer perceptron (MLP) projection because mutational features are more reflective of cellular lineage, while Z_x_ is mapped through a linear projection layer. The resulting projected representations, which may carry different weights or emphasis for each modality, are then concatenated along the feature dimension and fed into a bidirectional long short-term memory (BiLSTM) network to model cellular dynamics and predict mutational–transcriptional states across generations. The BiLSTM outputs a fused hidden representation at each time step, encoding both historical and future contextual information from the joint mutational-transcriptional state. Next, a learnable Weight Net module generates dynamic weights, enabling the splitting of the fused representation into modality-specific components. These components are then mapped through linear projection heads to produce the final predicted transcriptional states and mutational states at each generation, from which the corresponding generation N and expression matrix are inferred using scMut and scVI. MutTracer is able to infer both ancestral (t−1) and future (t+1) cellular states, providing a temporally resolved reconstruction of cellular lineage dynamics that integrates mutational and transcriptional information. By leveraging CGI as a temporal scaffold, MutTracer enables generation-aware inference of both ancestral and future transcriptional states, bridging measured cells and unobserved lineage stages.

### Mutations detected in scRNA-seq contain lineage information

To assess whether somatic mutations detected from scRNA-seq data encode meaningful lineage information, we systematically analyzed multiple tumor and lineage-tracing datasets.

First, we evaluated the reliability of mutation detection from scRNA-seq by comparing scRNA-seq and matched whole-genome sequencing (WGS) data from a cutaneous squamous cell carcinoma (cSCC) sample(Wiens et al., 2024). After quality filtering, approximately 60% of mutations identified in scRNA-seq were also detected in WGS (Fig. 2A), indicating that a substantial fraction of scRNA-seq–derived mutations reflects true underlying genomic variation.

**Figure 2.**
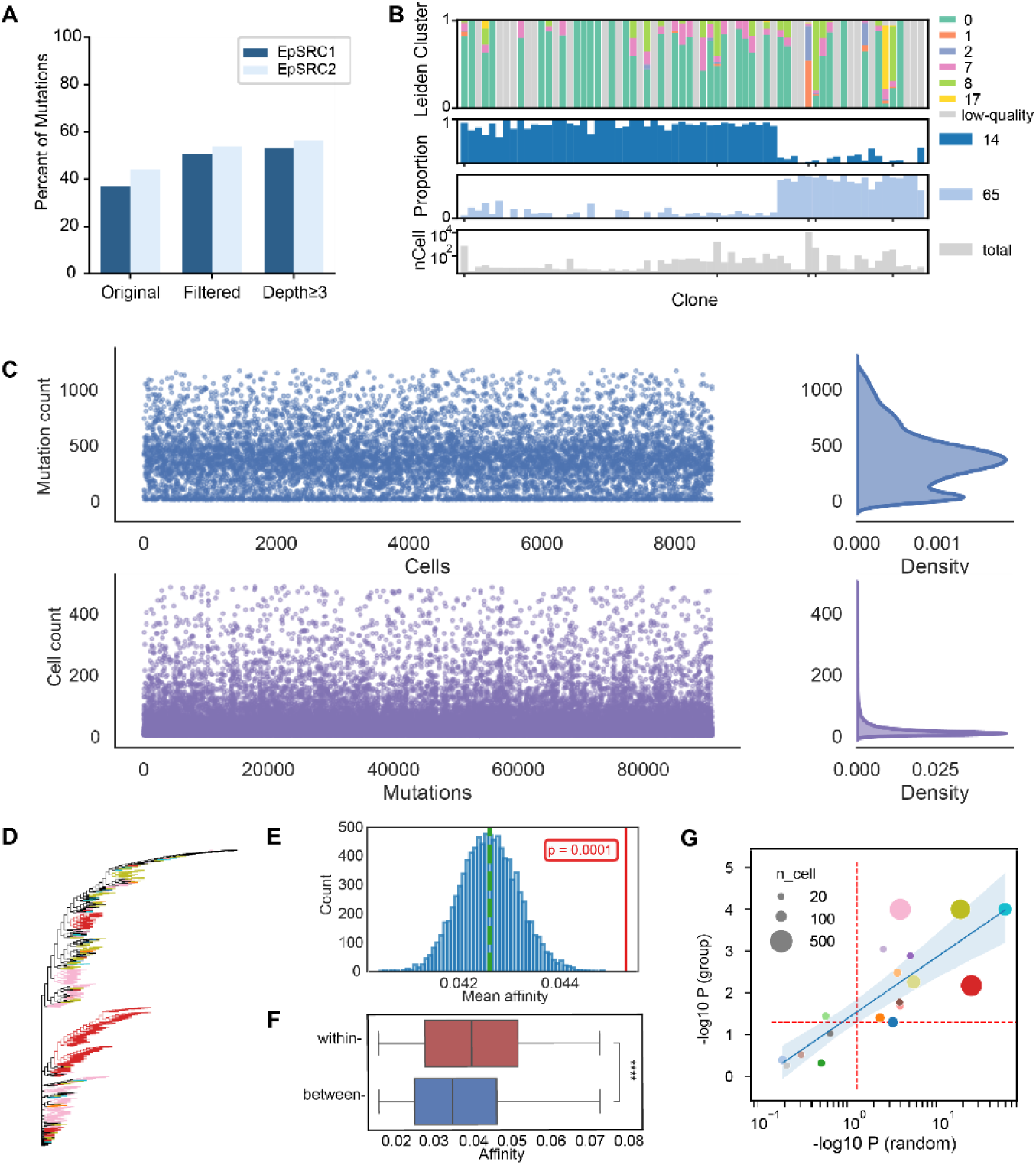
Mutations detected in scRNA-seq of tumors capture lineage information. (A) Percentage of mutations detected from cSCC scRNA-seq data at different filtering levels. (B) Cells are ordered by inferred clone identity along the x-axis. Top, Leiden cluster assignments for individual cells. Middle panels, proportions of cells assigned to each clone stratified by quality category. Bottom, total number of cells per clone. (C) Overview of mutation sparsity in the M100k mouse xenograft dataset. Top, scatter plot showing the number of detected mutations per cell across all cells. Bottom, scatter plot showing the number of cells in which each mutation is detected with the corresponding marginal density distributions. (D) Phylogenetic tree reconstructed from the mutation matrix corresponding to 19 barcode-defined clonal populations, where cells are colored by clone identity. (E) Null distribution of mean clonal affinity values obtained by random permutation of clone labels. The observed mean affinity of the barcode-defined clones is indicated by the red vertical line. (F) Comparison of phylogenetic affinity within individual barcode-defined clones versus between cells from different clones. Within-clone affinity were consistently smaller than between-clone affinity. (G) Significance of clone-specific clustering in the scRNA-seq mutation-based phylogenetic tree. The x-axis shows −log10 P-values from random permutation tests and the y-axis shows −log10 P-values for the comparison of within-clone versus between-clone phylogenetic distances. Point size reflects clone size.

To further assess whether mutations detected from scRNA-seq encode lineage-related information, we analyzed a CRISPR-based single-cell lineage tracing dataset of metastatic pancreatic cancer(Simeonov et al., 2021). Using VireoSNP (Huang et al., 2019), cells were assigned to clones based on the mutation matrix (M) derived from scRNA-seq data. The resulting mutational patterns were consistent with the clonal structure defined by CRISPR target sites, whereas clustering based solely on gene expression was insufficient to resolve these clones (Fig.2B). This result demonstrates that mutations detected from scRNA-seq can recover lineage structure that is not apparent from transcriptional similarity alone. Consistently, based on the theory of sparse correlations in MethylTree(Chen et al., 2025), a correlation analysis of mutations from lung adenosquamous cell carcinoma (LUAS) single-cell dataset(Qin et al., 2024) revealed distinct lineage structures corresponding to adenocarcinoma (ADC) and squamous cell carcinoma (SCC) (Fig.S1A). Together, these results demonstrate that mutations detected in tumor scRNA-seq data contain informative lineage signals, while their sparsity and noise preclude reliable use as discrete lineage markers, motivating alternative strategies that extract cumulative and continuous lineage information

Next, we extended our analysis to a lineage tracing scRNA-seq dataset from a mouse xenograft model of metastatic lung cancer(Quinn et al., 2021). In the M100k dataset, we extracted a mutation matrix containing 295,424 cells and 300,053 mutations after filtering for downstream clonal analysis (Fig.2C). We further filtered the mutation matrix to retain only the 19 barcode-based clonal populations reported in the original study, comprising a total of 2,043 cells. A phylogenetic tree was first reconstructed from scRNA-seq mutation profiles using all retained cells. We then quantified the degree of clonal aggregation on the tree by comparing the observed within-clone phylogenetic affinity to a null distribution generated by random permutation of clone labels across the fixed tree topology. Barcode-defined clones exhibited significantly stronger clustering than expected by chance (p = 0.0001) (Fig. 2D–F), indicating preferential aggregation of clonally related cells on the phylogenetic tree. Clone-specific phylogenetic analyses further showed that 15 of the 19 clones exhibited significant aggregation within the global mutation-based tree (P < 0.05) (Fig.2G, Fig.S1B), providing additional evidence that mutations detected in scRNA-seq capture lineage relationships.

Collectively, these results demonstrate that somatic mutations inferred from tumor scRNA-seq data are not randomly distributed but encode meaningful lineage information. However, the extreme sparsity and technical noise inherent to scRNA-seq mutation profiles limit their reliability as discrete lineage markers or for direct phylogenetic reconstruction. This motivates the development of alternative strategies that leverage the cumulative and continuous nature of mutation signals to infer cellular generation and temporal progression.

### Computational Strategies for Inferring Lineages from Sparse Mutation Data

Given that scRNA-seq mutations encode lineage information but are insufficient for stable tree-based reconstruction, we next consider algorithmic strategies for extracting robust cumulative temporal signals from sparse mutation data without explicit tree reconstruction. From an algorithmic perspective, existing lineage tree reconstruction methods based on dynamic labels typically assume a constant and continuous labeling rate, where the depth of a cell in the inferred tree reflects its lineage time. Our simulations demonstrate that this assumption is frequently violated in practice, although traditional phylogenetic methods can recover highly confident cell–cell relationships, they often yield unstable estimates of developmental progression (Fig.S2).

A critical solution lies in obtaining more robust and abundant mutational signals. High-throughput single-cell sequencing now enables the systematic use of somatic mutations as intrinsic lineage markers. Although somatic mutation data are often sparse and contain missing entries, the biological principle of progressive mutation accumulation generates a structured signal that can be exploited through mathematical modeling to reliably extract lineage features.

### Definition of Cellular Generation Index and Mutation Rate

To address the challenges posed by the high sparsity and false-positive rates of mutation matrices derived from scRNA-seq data, we developed scMut, a core component of VAETracer that leverages the CMM, which assumes that mutations accumulate progressively with cell divisions. Based on this principle, scMut jointly infers the CGI (N), a quantitative measure of lineage depth, and site-specific mutation rates (P) from the observed sparse mutation matrix (M). scMut further learns a low-dimensional mutational latent representation Z_m_, which captures underlying lineage structure. scMut integrates three complementary modules: gNMF for direct decomposition, VAE for probabilistic modeling and uncertainty quantification, and fine-tuning (FT) to refine parameter estimates using biological constraints. Together, N and P provide a low-dimensional, interpretable representation of lineage progression, where N captures the temporal order of cell divisions and P accounts for variable mutability across genomic sites.

To evaluate the model performance, we first generated simple simulated data under varying N and P, in which mutation states were assigned according to probabilities calculated from N and P. Results (Fig.S3, Table.1-2) show that both gNMF, VAE and FT can accurately recover the true N and P values even under a 10% noise mask.

To more realistically simulate biological lineage processes, we constructed a phylogenetic tree with a maximum depth of 50 generations, in which each cellular node inherits its mutation state from its parent; novel mutations are introduced with probability P only if the site remains unmutated in the parent, modeled via Bernoulli sampling. scMut performs robustly under these conditions, particularly at low mutation rates, which reflects the sparsity commonly observed in scRNA-seq-derived mutation matrices (Fig.3A, Fig.S4, Table.1-2).

**Figure 3.**
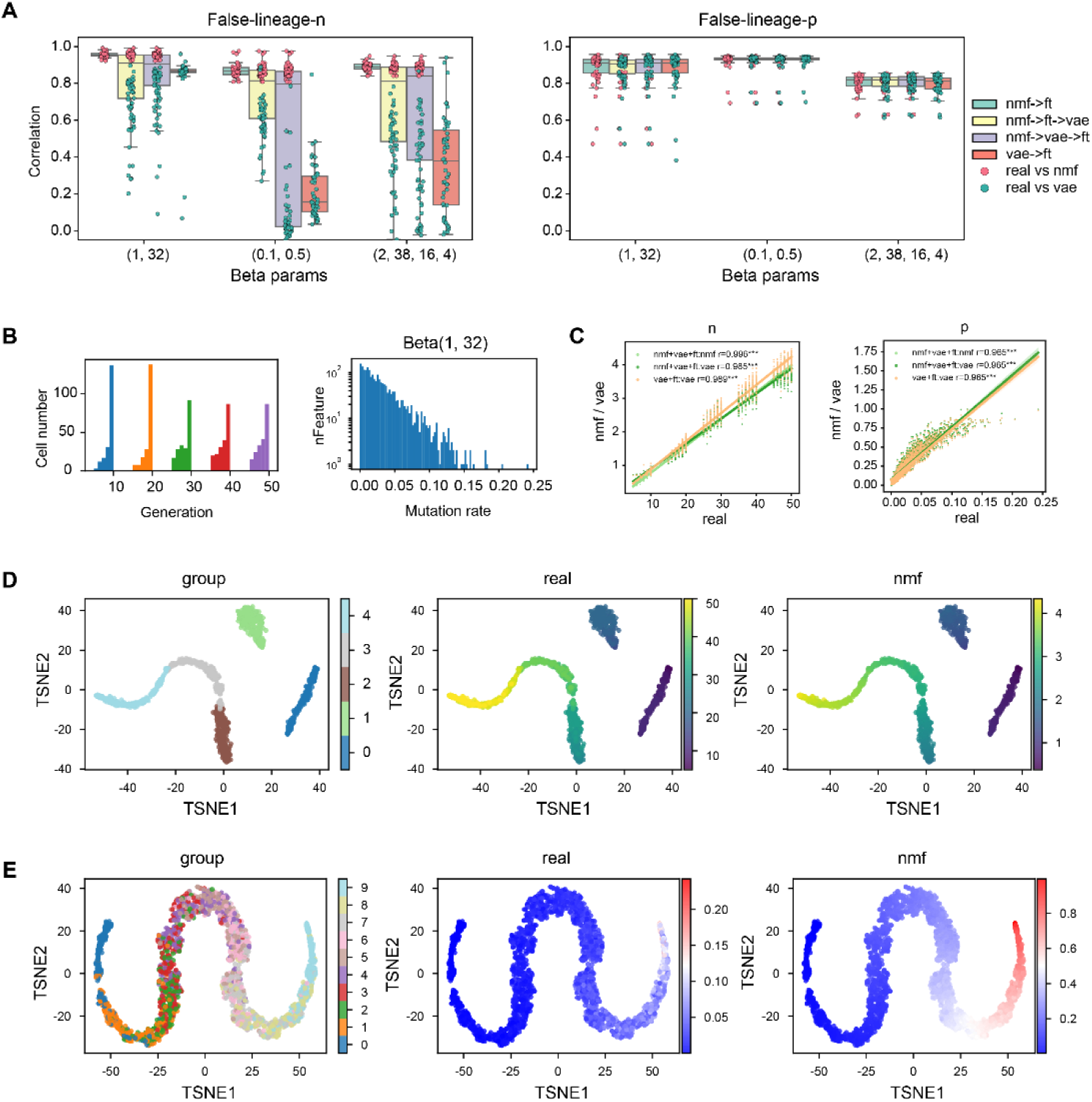
scMut models the mutational process parameters N and P. (A) Correlation between estimated and true N and P under varying mutation rates, using different module combinations in scMut. (B–C) Simulated temporal sampling and low mutation rate conditions, showing recovered N and P from VAE and gNMF modules. (D–E) Latent space Z_m_ from VAE visualized by dimensionality reduction (t-SNE); group order for cells (D) or sites (E) is sorted increasingly, and color intensity in real and gNMF maps reflects value magnitude. All correlations are computed using Spearman’s method.

We further tested the model on a lineage dataset with discrete sampling time points and low mutation rates to emulate real experimental designs, and found that both N and P were accurately recovered (Fig.3B-C, Table.3). To assess sensitivity and consistency across data quality, we simulated scenarios with increasing levels of missingness, from 10% to 90%, and found that scMut consistently recovers highly concordant cellular generation estimates (Fig.S5). Meanwhile, both N and P exhibited progressive variation along the latent Z_m_, reflecting lineage-associated dynamics (Fig.3D-E). Collectively, these results demonstrate the robustness of scMut under conditions of sparse mutational signals and its applicability to lineage evolution analysis.

Moreover, we show that the inferred site-specific P improves the accuracy of conventional phylogenetic tree reconstruction (Fig.S6), particularly in datasets with highly heterogeneous mutation origins, where uniform rate assumptions fail.

In summary, the N provides a quantitative measure of developmental time, while joint modeling of P improves inference accuracy by accounting for heterogeneous mutability. By leveraging consistent mutational patterns across cells (for P) and sites (for N), the scMut model robustly captures lineage signals even under high noise and extreme sparsity. Critically, the model bypasses phylogenetic tree reconstruction and directly infers generational relationships from the mutation matrix, enabling scalable and accurate lineage analysis without reliance on error-prone tree topologies.

### Application of scMut to tumor single-cell datasets

We next applied scMut to real tumor single-cell datasets to evaluate its ability to resolve tumor evolutionary progression. Based on the simulation results showing superior accuracy and robustness of gNMF in estimating N and P, we used the gNMF module to infer these two parameters in real data. The VAE module was used to learn the latent representation Z_m_. All analyses in this section were based on this workflow.

For the M100k dataset, we applied phylogeny-aware imputation to address high missingness in the single-clone mutation matrix. The strong correlation between N values before and after imputation indicates that lineage structure is preserved (Fig.4A), supporting reliable comparison with tree-based methods, which are sensitive to missing data. Phylogenetic trees were reconstructed for each single-clone mutation matrix, and mutational clusters, representing subclones, were identified and visualized using the VAE-derived latent space Z_m_. Taking clone-13 as an example, both N and clustering recapitulate the phylogenetic topology (Fig.4B-C). Across clones, N shows high concordance with node depth in the tree (Fig.4D), demonstrating its accuracy as a proxy for evolutionary progression. Furthermore, clusters in Z_m_ align closely with subclades from the phylogenetic tree (Fig. 4E), indicating that Z_m_ indeed captures biologically meaningful lineage relationships and subclonal structure. We also show that by combining inferred N values with phylogenetic topology, we can estimate N at internal nodes and assign branch lengths in units of cellular generations, enabling a quantitative, time-resolved lineage tree that is not achievable with conventional tree-building methods (Fig.S7). Together, these results show that scMut recovers robust lineage information without relying on explicit tree reconstruction.

**Figure 4.**
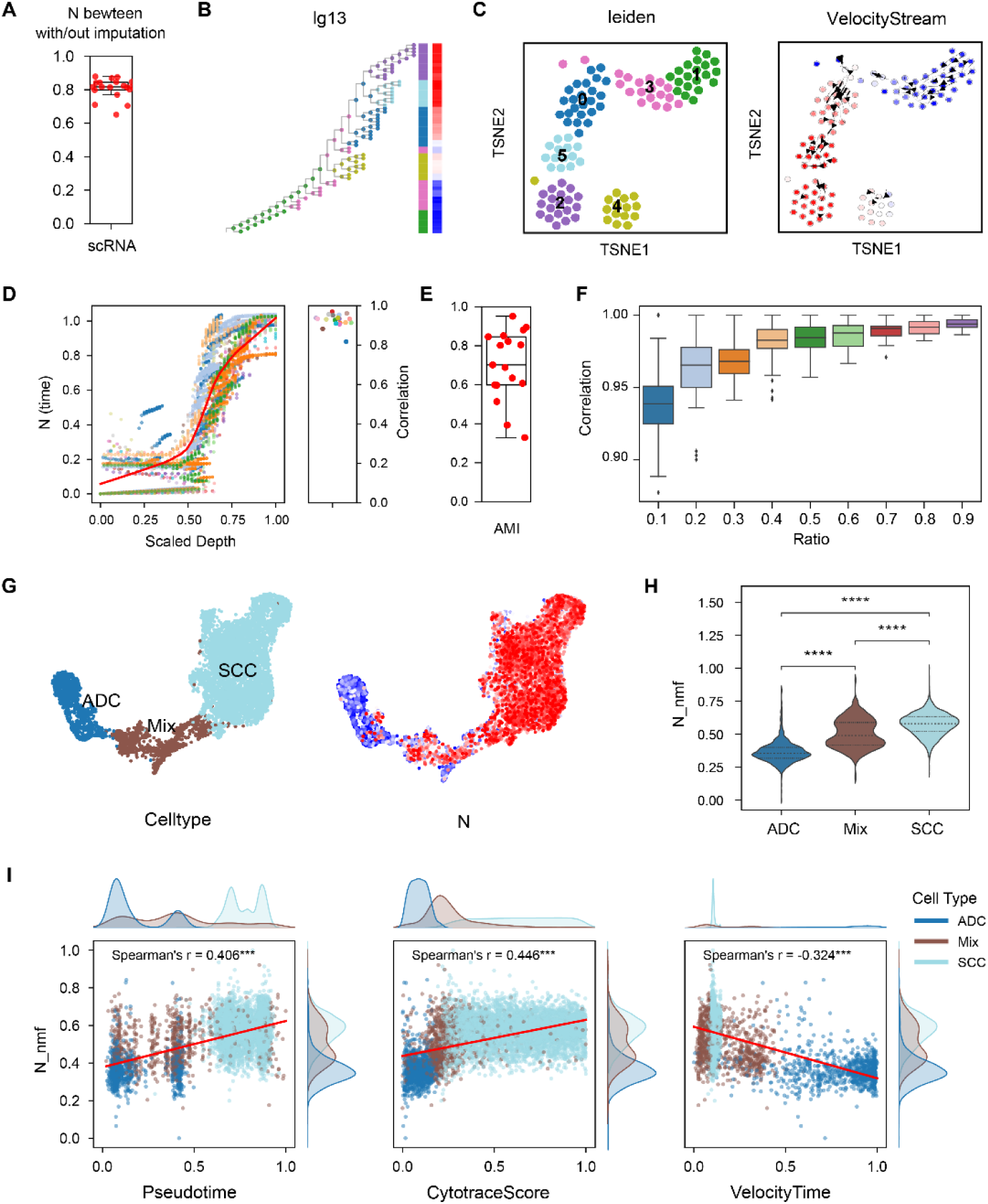
scMut applied to tumor single-cell datasets. (A) Comparison of N values before and after phylogeny-aware imputation. (B–C) Clone-13 analysis (here denoted as lg13): (B) bar plots showing Leiden clusters from Z_m_ (left) and distribution of N (right), colored to match the two subplots in (C); (C) t-SNE visualization of Clone-13 maps with left subplot showing cluster assignment and right subplot showing N, where arrows indicate lineage transmission direction from the phylogenetic tree. (D) Relationship between inferred N and scaled phylogenetic node depth across clones, where nodes (leaf) and clones (right) are colored by clone identity. (E) Z_m_ clusters compared with phylogenetic subclades, with cluster similarity measured by adjusted mutual information (AMI). (F) N consistency of Clone-13 under downsampling, shown as correlation at different sampling ratios. (G–I) Analysis of LUAS dataset: (G) spatial mapping of N, (H) group comparison of N by cell type, (I) comparison of N with pseudotime scores from Monocle, CytoTRACE, and RNA velocity. All correlations are computed using Spearman’s method.

To evaluate the robustness of N inference, we performed downsampling analysis on scRNA-seq mutation matrices and observed highly consistent estimates (Fig.4F, FigS8A-C), demonstrating strong reproducibility under subsampling. We also applied the same scMut pipeline to Cas9-based editing data, and the results showed qualitatively consistent patterns with those from scRNA-seq data (Fig.S8D-F). This demonstrates that scMut is applicable not only to somatic mutation systems but also to any lineage recording technology that follows a cumulative mutation principle.

Furthermore, when applied to the LUAS dataset, the model accurately recapitulated cellular generations, capturing the progression from ADC to SCC in agreement with previously reported trends(Tang et al., 2023) (Fig.4G). The inferred N showed significant differences among the three cellular states consistent (Fig.4H). When compared with established pseudotime methods, including Monocle (Trapnell et al., 2014), CytoTRACE (Gulati et al., 2020), and RNA velocity (La Manno et al., 2018), the ordering of cells was broadly concordant, but scMut revealed a more continuous and biologically plausible trajectory, reflecting the gradual and overlapping nature of cell state transitions during differentiation (Fig.4I).

In parallel, analysis of site-specific P revealed genes and pathways with significantly higher or lower mutational burden. Functional annotation of these genes provides a complementary, mutation-centric view of lineage-associated molecular programs (Fig.S9).

Collectively, these results indicate that scMut effectively captures lineage information from single-cell data, with the inferred CGI representing the temporal progression of cells and the latent features, offering a novel, mutation-derived perspective for delineating lineage-related cell populations.

### Integration of mutation-based lineage information for expression inference

Building upon the mutation-derived lineage representations and CGI-inferred cellular generations from scMut, we next investigated how lineage information can be leveraged to improve time-resolved inference of gene expression dynamics. Although scRNA-seq captures only a snapshot of cellular states, cells measured within a single experiment can originate from multiple cellular generations, reflecting lineage heterogeneity at the time of sampling. These observed generational states provide critical anchors for modeling temporal progression and enable the inference of ancestral or descendant cellular states that are not directly measured.

While transcriptional similarity alone is insufficient for reliable inference of directional state transitions across discrete time points, particularly when extrapolating to unobserved intermediate or ancestral states, somatic mutations accumulate irreversibly along cell divisions and thus provide a more stable signal of temporal ordering. To incorporate such lineage-derived temporal constraints, we integrated mutation-informed latent variables with transcriptional embeddings to guide the modeling of gene expression dynamics. Specifically, the mutational latent representation Z_m_, which encodes cumulative lineage information, and the transcriptional embedding Z_x_ were jointly projected into a shared latent space and modeled using a bidirectional recurrent architecture. By assigning greater emphasis to Z_m_ in this integration, the model is guided by lineage-consistent temporal structure while retaining transcriptional variability, thereby enabling the reconstruction of unmeasured cellular states across generations.

To evaluate whether mutation-guided temporal modeling improves inference of unobserved intermediate states, we applied MutTracer to simulated datasets with explicitly defined cellular generations. Using the same simulation framework described above, we generated datasets with three discrete time points (Time 1–3). Mutation matrices were simulated to derive N and Z_m_ (Fig.5A), while gene expression matrices were generated using dyngen (Cannoodt et al., 2021) to model linear transcriptional dynamics (Fig.5B). Expression representations (Zₓ) were obtained using scVI (Lopez et al., 2018), and cells were grouped into the same three time points based on the simulated expression dynamics. Using joint Zₘ–Zₓ representations from Time 1 and Time 3, we predicted gene expression at Time 2 and compared the results with LineageVAE (Majima et al., 2024) and WOT (Schiebinger et al., 2019) (Fig.5C). Our model produced expression representations that were closer to the real Zₓ_₂_ (Fig.5D), demonstrating improved accuracy in reconstructing intermediate states along cellular evolutionary trajectories. In addition, the predicted Zₘ_2_ and inferred N were closer to the distribution of real Time 2 (Wasserstein distance = 0.416) (Fig.5E-F). Forward and backward inference of N across time points produced distributions closely aligned with the ground truth (Fig.S10B). Consistent trends were observed for the inferred adjacent latent states, including Z_mt±1_ and Z_xt±1_, which also closely matched their corresponding ground truth distributions.

**Figure 5.**
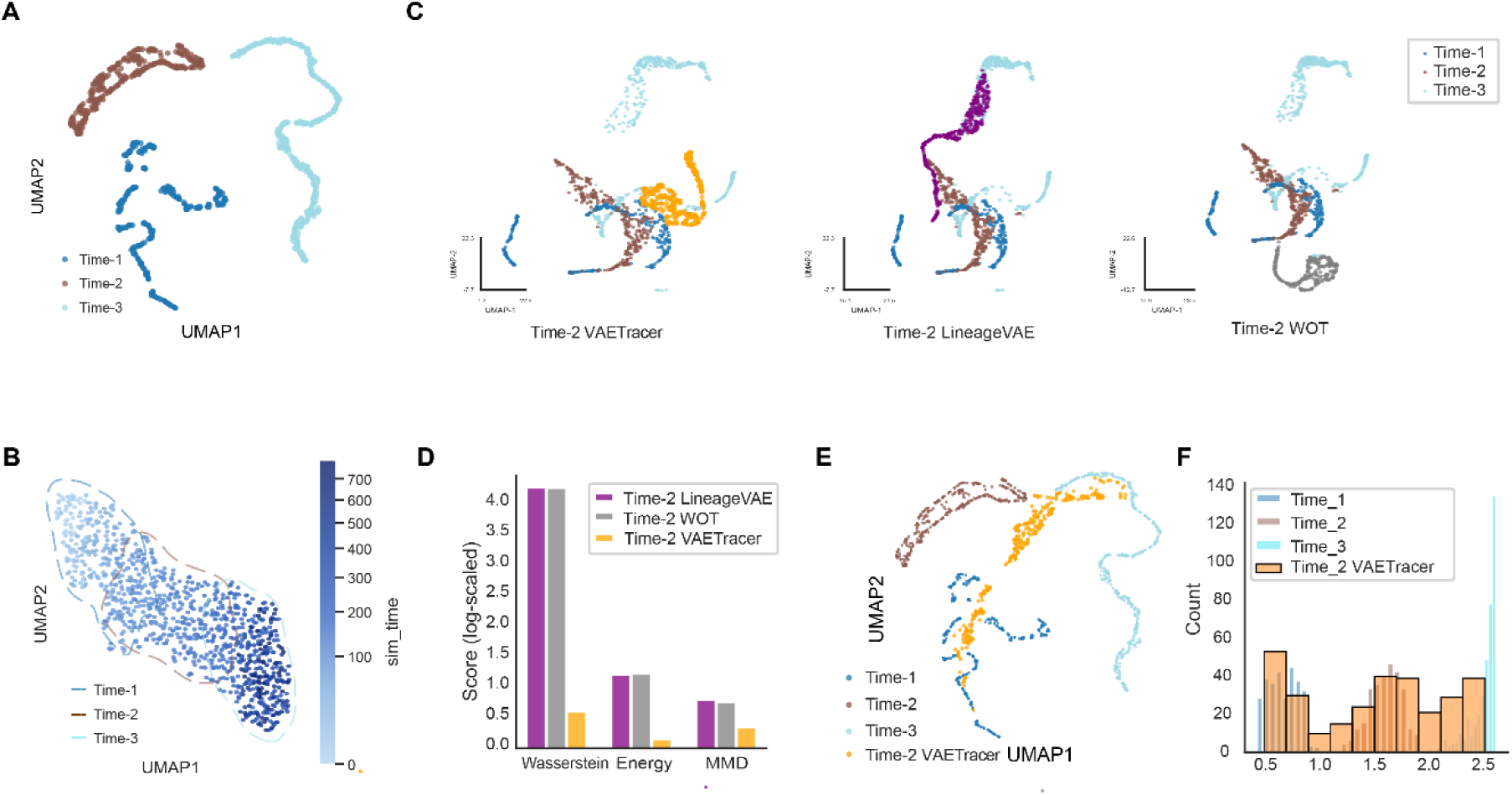
Integration of mutation-based lineage information improves time-resolved inference of gene expression dynamics. (A) UMAP visualization of simulated mutational latent representations derived from scMut. Cells are colored by discrete generational time points (Time 1–3). (B) UMAP visualization of simulated transcriptional states. Points are colored by simulated continuous time (sim_time), with contours indicating Time 1–3. (C) UMAP projections of predicted intermediate (Time 2) expression states generated by different methods. From left to right: VAETracer, LineageVAE, and WOT. Predicted Time 2 cells are overlaid on the real Time 1-3 cells for comparison. (D) Quantitative comparison of predicted Time 2 expression representations with the real Time 2 distribution using Wasserstein distance, energy distance and maximum mean discrepancy (MMD). (E) UMAP visualization of predicted Time 2 mutational states from VAETracer overlaid with real cells from Time 1–3. (F) Distribution of N for real Time 1–3 cells and predicted Time 2 cells from VAETracer.

Together, these results suggest that the model effectively predicts joint mutational and expression features across time points, capturing both forward and backward dynamics of coupled mutational and transcriptional states along cellular evolutionary trajectories. These findings also motivate its application to real tumor datasets with complex generational heterogeneity.

### Application of MutTracer to tumor single-cell datasets

Having established the accuracy of MutTracer on simulated datasets, we next applied the model to the LUAS tumor single-cell dataset to evaluate its ability to reconstruct temporal expression dynamics in a biologically complex system, in which ADC, mixed (Mix), and SCC populations represent progressive tumor states, and assigned them to discrete time points 1–3 to evaluate the model’s bidirectional prediction performance.

Predictions of Z_xt±1_ were compared with LineageVAE and WOT across forward, intermediate, and backward inference tasks (Fig.6A-C). The predicted Zₓ representations were decoded into expression matrices (X_t±1_) using the scVI decoder, and R² with the observed marker expression at the corresponding time points showed strong agreement (Fig.6D-F), confirming the model’s accuracy in capturing temporally resolved expression dynamics from mutation-informed features. In parallel, the predicted Zₘ representations were encoded to CGI (N_t±1_) using the scMut encoder N, enabling direct evaluation of generational progression in predicted states (Fig. S11A).

**Figure 6.**
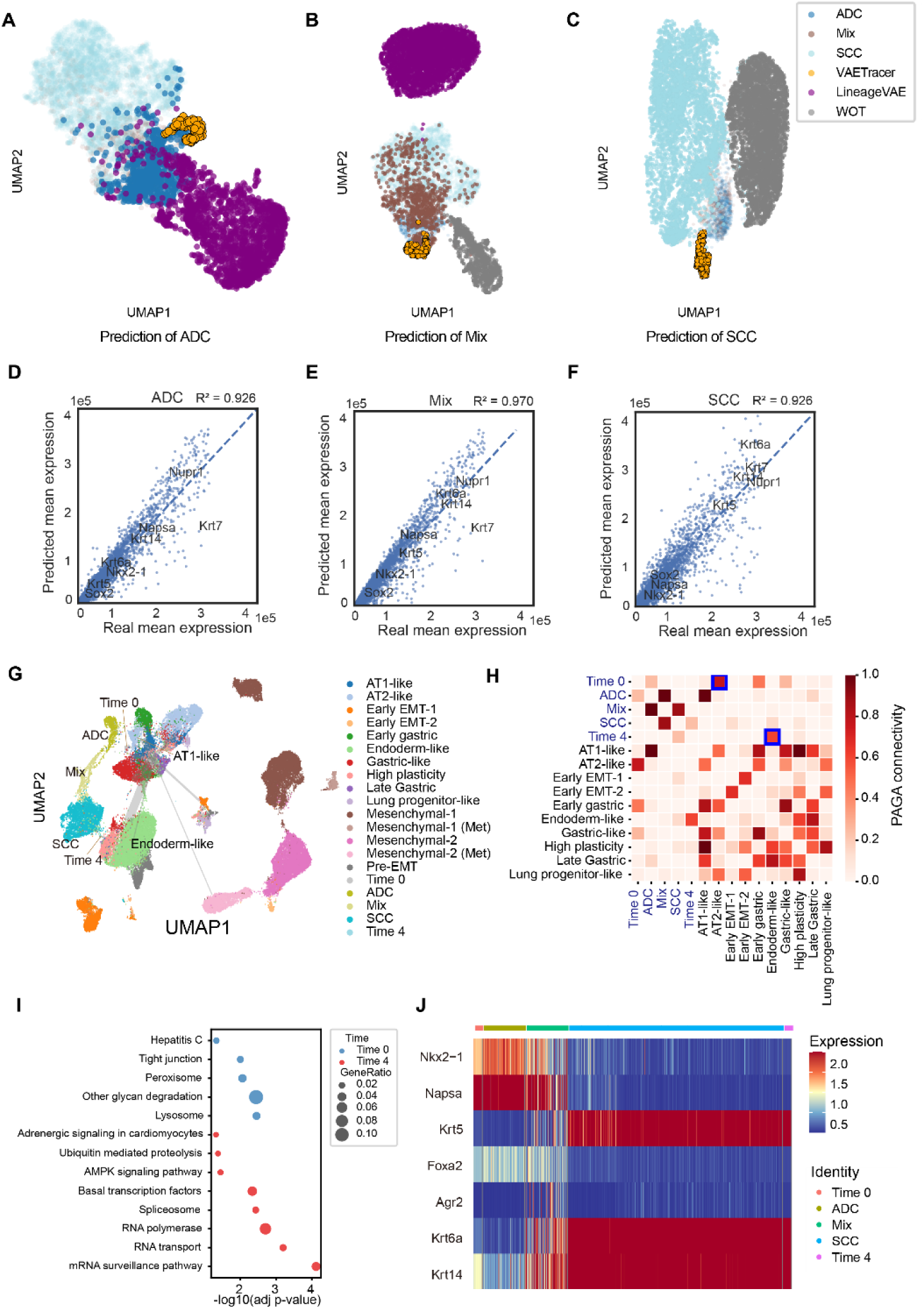
Application of MutTracer to the LUAS single-cell tumor dataset. (A-C) UMAP visualization of inferred Zₓ for LUAS tumor cells under three prediction tasks. A, Forward prediction of ADC states. B, Intermediate prediction of mixed (Mix) states. C, Backward prediction of SCC states. (D-F) Comparison of predicted and observed mean expression levels of marker genes for each tumor subtype. Each point represents a gene and dashed lines indicate linear regression fits. D, ADC prediction. E, Mix prediction. F, SCC prediction. (G) PAGA-based UMAP visualization integrating the Kras;Trp53-driven lung adenocarcinoma dataset, the LUAS dataset, and the predicted Time 0 and Time 4 expression states inferred by MutTracer. (H) PAGA connectivity matrix showing similarity between predicted Time0 and Time4 states and reference lung epithelial cell populations. (I) Gene set enrichment analysis of marker genes identified from predicted Time 0 and Time 4 expression profiles. (J) Heatmap of normalized expression for selected marker genes across predicted temporal states (Time 0, Time 4) and observed LUAS subtypes (ADC, Mix, SCC).

To further explore early- and late-stage tumor states, all LUAS cells were input into the model to predict Time 0 (earlier stage) and Time 4 (later stage) expression profiles. Comparison with lineage-tracing Kras;Trp53-driven lung adenocarcinoma dataset (Yang et al., 2022) revealed that the predicted Time 0 state closely resembled the AT2-like cell program, which is associated with progenitor-like functions in early tumorigenesis (Li et al., 2025a; Wang et al., 2021). In contrast, the predicted Time 4 state was more similar to endoderm-like cells (Fig.6G-H), consistent with the later-stage, more differentiated states observed during tumor progression (Li et al., 2025a). Consistently, visualization of the Z_m_ₜ revealed an ordered progression from predicted Time 0 to Time 4, further supporting the coherence of the inferred temporal trajectory (Fig. S11B). Functional enrichment analysis revealed that Time 0 marker genes were enriched for pathways involved in cellular homeostasis and metabolic regulation (Wang et al., 2025), reflecting the maintenance of progenitor-like states, whereas Time4 marker genes showed enrichment in pathways related to transcriptional regulation, RNA processing, and signaling (Wang et al., 2021) (Fig.6I). In addition, the expression dynamics of known LUAS, ADC, and SCC marker genes across Time 0-4 recapitulated expected temporal trends (Fig.6J), highlighting the ability of MutTracer to capture biologically meaningful, mutation-informed temporal progression of tumor states.

## Discussion

In this work, we present VAETracer, a mutation-informed framework that enables the reconstruction of cellular lineage structure and temporal transcriptional dynamics directly from scRNA-seq data. By leveraging the cumulative and irreversible nature of somatic mutations, VAETracer provides an internal temporal reference that is independent of transcriptional similarity, addressing a fundamental limitation of expression-based trajectory inference methods in heterogeneous tumor systems.

Although scRNA-seq experiments capture cells at a single experimental time point, the sampled population often comprises cells originating from multiple cellular generations. This latent generational heterogeneity reflects ongoing lineage evolution at the time of sampling but is not explicitly observable from transcriptional profiles alone. By extracting a mutation-derived CGI, VAETracer reveals this hidden temporal structure and uses it to anchor the modeling of gene expression dynamics, enabling inference of ancestral or descendant cellular states that are not directly measured.

Unlike classical lineage reconstruction methods, VAETracer does not rely on accurate tree topologies, which are difficult to obtain from noisy scRNA-seq–derived mutation data. At the same time, it extends beyond conventional pseudotime methods by introducing mutation-informed temporal constraints, resulting in more biologically plausible and directionally coherent state transitions.

The framework is inherently extensible to other data modalities that follow a cumulative inheritance pattern, including artificial lineage barcodes, mitochondrial mutations, and epigenetic alterations. The core principle of leveraging cumulatively inherited mutational signals to infer lineage and temporal progression is not restricted to scRNA-seq data and can, in principle, be extended to chromatin accessibility or genomic mutation profiles, such as single-cell ATAC-seq(Buenrostro et al., 2015) or single-cell Whole-Genome Sequencing(Qiao et al., 2025). Moreover, the increasing availability of multimodal and spatially resolved single-cell datasets provides opportunities to integrate mutation-informed lineage signals with regulatory or spatial information(Stahl et al., 2016), enabling joint analysis of temporal, molecular, and spatial dimensions of tumor evolution. Beyond cancer, VAETracer may also be applicable to developmental, regenerative, or aging systems in which somatic mutations accumulate over time, particularly as advances in single-cell sequencing technologies enable more sensitive and accurate detection of somatic variants in non-malignant tissues.

Despite its advantages, the performance of VAETracer remains dependent on the quality of mutation detection from scRNA-seq data. Current protocols are limited by sparse coverage and technical noise, particularly when mutations are primarily detected in the 3′ untranslated region, resulting in incomplete and noisy mutation matrices. Continued improvements in single-cell sequencing technologies and probabilistic variant calling strategies are expected to further enhance the robustness and applicability of mutation-informed lineage modeling.

In summary, VAETracer offers a scalable and integrative approach for uncovering lineage structure and temporal transcriptional dynamics from widely available scRNA-seq data. By reframing somatic mutations as a generational temporal signal rather than a basis for explicit phylogenetic reconstruction, VAETracer provides a practical framework for time-resolved analysis of cellular evolution in complex biological systems.

## Methods

### Module Pre-process: variant calling from scRNA-seq data and phylogenetic reconstruction

#### 1.1 Mutation calling from scRNA-seq data

The structure of this module is shown in Fig.1A. Single-cell RNA-seq reads were aligned to species-matched reference genomes using STAR (v2.6.0a), including GRCm38.p6 for mouse samples and GRCh38.p13 for human samples. Gene expression quantification and BAM file generation were performed separately using CellRanger (v7.0.1), and the resulting BAM files were used for downstream variant calling analyses. Aligned reads were sorted and indexed using SAMtools (v1.9), and read group information was added using Picard AddOrReplaceReadGroups (v2.18.29). PCR duplicates were marked and removed using picard MarkDuplicates. Base quality score recalibration (BQSR) was performed using GATK (v4.2.3.0)(McKenna et al., 2010) BaseRecalibrator and ApplyBQSR with known variant sites provided as references. To enable variant calling from spliced RNA reads, SplitNCigarReads was applied following GATK best practices for RNA-seq data. Processed BAM files were restricted to standard autosomes and sex chromosomes prior to variant calling. Variants were called for each sample using GATK HaplotypeCaller in GVCF mode. Per-sample GVCFs were jointly genotyped across all samples using GenomicsDBImport followed by GenotypeGVCFs on a per-chromosome basis. Chromosome-level VCFs were subsequently merged into a single VCF file for downstream analysis, and cell-level mutation matrices were generated by extracting allele information from scRNA BAM files using a custom script (GetAF.py).

#### 1.2 Phylogenetic Reconstruction from scRNA-seq mutations

Mutations were further filtered using custom minimal thresholds on the number of supporting cells and mutations before reconstructing phylogenetic trees with Cassiopeia (v2.1.0)(Jones et al., 2020) using the Neighbor-Joining algorithm.

### Module scMut: Joint Inference of Cellular Generations and Mutation Rates via Probabilistic Latent Modeling

The structure of this module (scMut) is shown in Fig.1B. scMut is a probabilistic model based on the sampling process of mutation accumulation, designed to jointly infer the N and site-specific P from an observed mutation matrix M with values in [0, 1, 3], where rows represent cells, columns represent genomic sites, and entries indicate wild-type (0), mutated (1), or missing (3) states. The CMM assumes that mutations are acquired independently across sites and accumulate monotonically with lineage depth. This enables the disentanglement of N and P from the observed sparse M, despite technical noise and dropout. Given the binary M is treated as Bernoulli realizations of the underlying probability matrix R, the framework supports inputs in either binary or probabilistic form. The details of this module can be described in the following sections, i.e. Sections 2.1-2.6.

#### 2.1 Cumulative mutation model

We introduce the cumulative mutation model (CMM), a probabilistic framework that describes how somatic mutations accumulate irreversibly across cell divisions. The model assumes that at each cell division, a genomic site j has a fixed probability p_j_∈(0,1) of acquiring a novel mutation, and once a mutation occurs, it is stably inherited by all descendant cells.

For a cell i with generational index n_i_, the probability that site j remains unmutated after n_i_ independent divisions is (1 − p_j_)^ni^. Therefore, the probability r_ij_ of observing a mutation is:

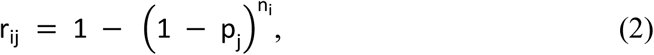

This formulation implies that r_ij_ increases monotonically with both n_i_ and p_j_. Under the CMM, the observed mutation m_ij_ ∈{0,1} is generated by independent Bernoulli trials:

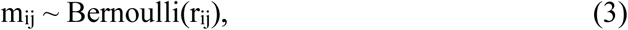

Thus, the observed mutation matrix M = [*m*_*ij*_] can be viewed as a noisy realization of the underlying probability matrix R = [*r*_*ij*_], enabling flexible input representation, either binary or probabilistic. For simplicity, we use M to represent the probability R consistently in Fig.1 and the main text, as defined in Eq. 1.

#### 2.2 Silicon-data simulation

Mutation rates (P) across genomic sites were simulated using a beta distribution to capture heterogeneous mutability. Specifically, site-specific P were drawn from either a unimodal Beta (a, b) or a bimodal mixture of two beta distributions, enabling flexible modeling of diverse mutation rate profiles. Given fixed N and P, we compute the observed mutation matrix M under the CMM, followed by introduction of missing entries according to a predefined noise level.

To simulate realistic lineage structures, a binary branching process was used to grow a cell lineage tree from a single root. Each cell divides stochastically, with a survival rate controlling the probability of continued division. Mutations are accumulated irreversibly during each division. Terminal cells were sampled from specified generation intervals to control age distribution, and their mutation profiles were used to construct the final dataset along with an associated Newick-formatted phylogenetic tree.

#### 2.3 Non-negative Matrix Factorization for parameter decomposition

As discussed in the main text, the cumulative mutation model (CMM) shares a structural analogy with NMF, reducing to the standard M=NP form under specific conditions (such as low P) by linearizing the right-hand-side of eqn.(1). To solve the full CMM without linear approximation, we adopt a generalized NMF (gNMF) framework that directly models the underlying generative process. The objective is defined as:

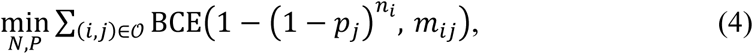

where BCE denotes binary cross-entropy loss, and the sum is restricted to non-missing entries (*m*_*ij*_≠ 3). Optimization is implemented using PyTorch with Adam gradient descent.

#### 2.4 Variational Autoencoder Framework for Probabilistic Inference

The VAE framework is designed to operate in two distinct modes, enabling flexible parameter inference and representation learning. In mode 1 (’np’ mode), the architecture separates parameter estimation from reconstruction. It consists of:

i. An encoder that maps M to a latent *Z*_*m*_;
ii. A decoder that further compresses *Z*_*m*_ into N (via softplus transformation);
iii. A globally shared parameter P, represented as trainable logits and transformed via sigmoid.

Together, N and P are used to compute the expected mutation probability matrix *R̂* through the CMM model, which is then compared to M.

In mode 2 (‘xhat’ mode), the model adopts a standard symmetric VAE structure: an encoder maps input M to latent space *Z*_*m*_, and a decoder reconstructs *M̂* directly.

Training in this work was conducted exclusively in ‘np’ mode. While ‘xhat’ mode can theoretically yield latent representations similar to those from ‘np’ mode, the latter provides direct estimates of biological parameters and thus offers greater interpretability. Notably, *R̂* in ‘np’ mode is theoretically constrained by the outer product structure of N and P, which may lead to underfitting, whereas *M̂* in ‘xhat’ mode is not subject to such constraints.

#### 2.5 Parameter-Specific Loss Functions and Training Strategy

In ‘np’ mode, the loss function measures reconstruction accuracy based on the predicted mutation probabilities:

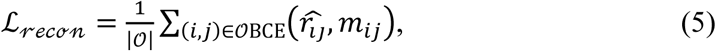

where 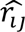 is computed from the current estimates of *n*_*i*_ and *p*_*i*_.

A KL divergence term that regularizes the latent distribution against a standard normal prior:

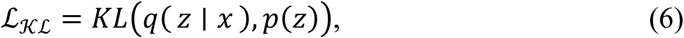

If prior estimates (e.g., from gNMF) are available, they are incorporated via a mean squared error regularization term:

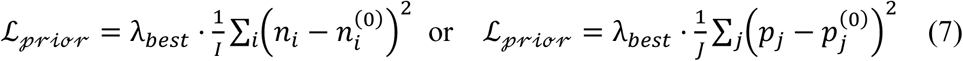

The total objective combines reconstruction and regularization:

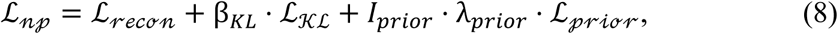

While in ‘xhat’ mode, the loss function of reconstruction becomes:

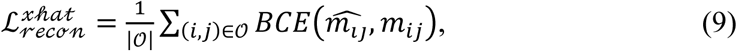

#### 2.6 Learning Site-Level Representations via Transposed Modeling

To learn low-dimensional representations of genomic sites (rather than cells), the input M is transposed such that sites become samples and cells become features. In this configuration:

i. The latent space *Z*_*m*_ encodes site-level mutational patterns,
ii. N becomes the globally shared parameter and P is compressed from *Z*_*m*_.

#### 2.7 Fine-Tuning with Biological Constraints

N values from gNMF or VAE are continuous and ordinally consistent but scale-dependent; if the true generation range is known, N and P can be fine-tuned for improved biological fidelity.

First, the continuous estimate of N is transformed into an integer-valued vector by applying an optimal linear scaling factor k. Specifically, we solve:

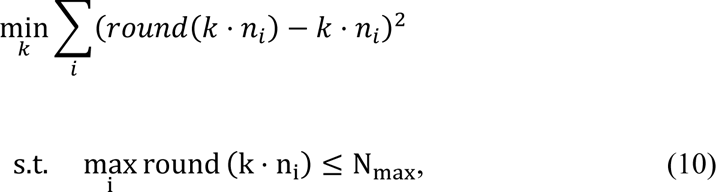

where *n_i_* denotes the inferred continuous generation index for cell i, and *N_max_* is a user-defined upper bound on the number of generations. The solution yields the optimal scaling factor *k* ∗, which, when applied to the *n_i_* and rounded, produces the fine-tuned integer N.

With N fixed, the P is then re-optimized by minimizing:

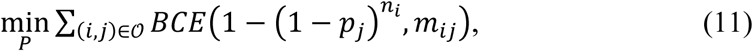

Only P is updated in this phase, effectively performing a constrained re-fitting analogous to gNMF. This ensures consistency between the final N and P, improving calibration of the generative model.

#### 2.8 Implementation and Optimization

All models are implemented in PyTorch. The Adam optimizer is used with a learning rate of 1e-3. Training uses mini-batches (ranging from 256 to 10,000), with early stopping based on validation loss (patience: 45 epochs). Numerical stability is maintained using log-space transformations and boundary clamping.

scMut outputs are processed using Scanpy and AnnData for downstream analysis and visualization (e.g., UMAP, tSNE).

### Module MutTracer: Integrative temporal inference of mutational and transcriptional states

The structure of this module is shown in Fig.1C. MutTracer is a generation-aware temporal modeling framework that jointly models mutational and transcriptional latent representations to predict cellular states across discrete generations. The model takes as input the mutational latent representation *Z*_*m*_ and the transcriptional embedding *Z*_*x*_, which together provide complementary views of lineage progression and cellular state.

Cells are first ordered and discretized into generational time points according to *N*. For each generation *t*, the corresponding mutational and transcriptional latent representations are integrated to construct a joint latent state, which is used to model transitions between neighboring generations. Temporal dependencies are captured using a bidirectional recurrent architecture, enabling prediction of latent states at adjacent generations in both forward and backward directions.

The detailed architecture of MutTracer consists of

i. joint projection of mutational and transcriptional latent features into a shared space;
ii. generation-aware bidirectional LSTM modeling of latent dynamics;
iii. supervised prediction of adjacent generational states with temporal regularization, as described below, i.e. sections 3.1-3.4.

#### 3.1 Joint latent projection of mutational and transcriptional features

For each cell at generation *t*, we obtained a transcriptional latent representation Z_x_t__ ∈ R^m_x_^ from scMut encoder Z and a mutational latent representation Z_m_t__ ∈ R^m_m_^ from scVI encoder. To facilitate unified temporal modeling while accounting for the heterogeneous characteristics of the two modalities, we apply modality-specific projection networks to transform the latent representations into a temporally compatible feature space. Specifically, the mutational representation is projected via a multi-layer perceptron (MLP),

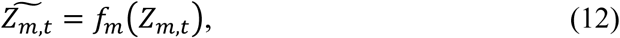

while the transcriptional representation is projected using a linear transformation,

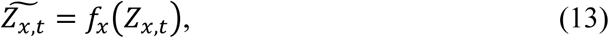

where *f*_*m*_(⋅) and *f*_*m*_(⋅) are learnable projection functions with different capacities, reflecting the higher sparsity and discreteness of mutational signals compared to transcriptional embeddings.

The projected features are then concatenated along the feature dimension to form a joint latent state

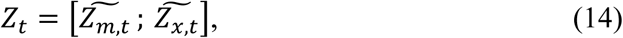

which serves as the input to the temporal dynamics model. This asymmetric projection strategy balances the representational contributions of mutational and transcriptional features and enables effective joint modeling of their temporal evolution.

#### 3.2 Bidirectional LSTM for generation-aware latent dynamics

To model generation-aware latent dynamics, we employ a bidirectional long short-term memory (BiLSTM) network as a temporal context encoder operating on the joint latent representations. For each cell, the projected mutational and transcriptional features are concatenated to form a sequence of joint latent states{*Z*_1_, *Z*_2_, …, *Z*_*T*_}ordered by cellular generation.

The concatenated latent sequence is processed by two independent LSTM networks in the forward and backward temporal directions. Specifically, the forward LSTM encodes contextual information from earlier generations, while the backward LSTM encodes information from later generations. The outputs of the two LSTMs are aligned in time and combined to produce a bidirectional hidden representation at each generation *t*:

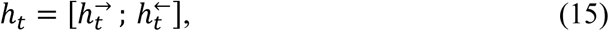

where 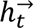 and 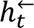 denote the forward and backward hidden states corresponding to generation *t*, respectively.

Importantly, although the BiLSTM is applied to the entire latent sequence during computation, the model is not trained to perform global sequence-to-sequence prediction. Instead, the hidden representation *h*_*t*_ serves as a generation-specific contextual embedding, summarizing information from neighboring generations surrounding *t*. This design allows the model to incorporate both ancestral and descendant context when modeling the latent state at each generation, without enforcing explicit long-range temporal dependencies.

The BiLSTM consists of two stacked layers with *H* hidden units per direction. The resulting bidirectional hidden representations provide a temporally smoothed and context-aware encoding that supports subsequent mutation- and transcription-specific predictions at each generation.

#### 3.3 Prediction of adjacent generational states and loss function

Given the joint latent representation at generation *t*, the concatenated mutational and transcriptional features are first processed by a BiLSTM network to capture temporal dependencies across generations. The forward and backward hidden states are concatenated to form a fused bidirectional representation *H*_*tt*_.

To decouple the fused temporal features into modality-specific components, a learnable weight network is applied to *H*_*tt*_, producing dynamic weights that modulate the contributions of mutational and transcriptional information. Conditioned on these weights, modality-specific prediction heads are used to infer latent states at adjacent generations:

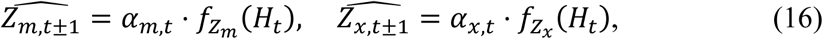

where *f_Z_m__*(⋅) and *f_Z_x__* (⋅) denote independent linear projection heads for the mutational and transcriptional latent representations, respectively, and α_*m,t*_, α*_x,t_* are modality-specific weights satisfying α_*m,t*_ + α_*x,t*_ = 1.

For generations with observed latent representations, the model is trained using a supervised reconstruction objective. Specifically, a mean squared error (MSE) loss is applied to both latent components:

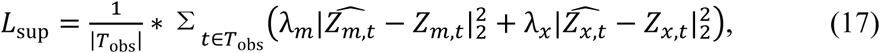

where *T*_obs_ denotes the set of generations with available observations.

The coefficients λ_*m*_ and λ*_x_* control the relative weighting of the mutational and transcriptional reconstruction errors. In practice, a higher weight λ_*m*_ is assigned to the mutational latent variable *Z*_*m*_ to emphasize accurate modeling of cellular state transitions across generations.

To ensure biologically plausible latent trajectories, additional temporal regularization terms are introduced, including constraints enforcing continuity between adjacent generations, smoothness based on higher-order temporal differences, and consistency with observed transition directions. The final training objective is defined as a weighted sum of the supervised reconstruction loss and the temporal regularization terms.

#### 3.4 Optimization and training procedure

Model parameters were optimized using the Adam optimizer with AMSGrad.

The learning rate was set to 1 × 10^−4^. Training was conducted for 200 epochs using mini-batches of size 30, with each mini-batch containing cells associated with a single barcode.

All models are implemented in PyTorch. MutTracer outputs were processed using Scanpy and AnnData for downstream analysis and visualization.

## Supporting information

Supplementary Tables and Figures VAETracer

## Acknowledgements

This work was supported by National Natural Science Foundation of China (T2341007, T2350003, 12131020, 42450084, 42450135, 12326614, 12426310, T2542018 and 62002329); National Key R&D Program of China (2022YFA1004800, 2025YFF1207900, 2025YFC3409300); Science and Technology Commission of Shanghai Municipality (23JS1401300); Zhejiang Province Vanguard Goose-Leading Initiative (2025C01114); Shenzhen Medical Research Fund (E250200621); JST Moonshot R&D (No. JPMJMS2021).

## Code availability

An open-source software implementation of VAETracer is available on Github (https://github.com/Kaiyu-W/VAETracer). The repository includes all scripts for module Pre-process, scMut, and MutTracer.

## Data availability

All single-cell RNA-seq and lineage tracing datasets analyzed in this study are publicly available. scRNA-seq and matched WGS data of cSCC sample were downloaded from GEO (GSE213338) and NCBI BioProject database (PRJNA1099909). The CRISPR-based single-cell lineage tracing dataset of metastatic pancreatic cancer were downloaded from GEO (GSE173958). The LUAS single-cell dataset was downloaded from the National Omics Data Encyclopedia (accession no. OEP003086). The mouse xenograft model of metastatic lung cancer was downloaded from GEO (GSE161363). The lineage-tracing Kras;Trp53-driven lung adenocarcinoma dataset were downloaded from NCBI BioProject database (PRJNA803321).

## Notes

### Competing Interest Statement

The authors have declared no competing interest.

## References

Anderson, D.J., Pauler, F.M., McKenna, A., Shendure, J., Hippenmeyer, S., and Horwitz, M.S. (2022). Simultaneous brain cell type and lineage determined by scRNA-seq reveals stereotyped cortical development. Cell Syst 13, 438–453 e435.

Andor, N., Lau, B.T., Catalanotti, C., Sathe, A., Kubit, M., Chen, J., Blaj, C., Cherry, A., Bangs, C.D., Grimes, S.M., et al. (2020). Joint single cell DNA-seq and RNA-seq of gastric cancer cell lines reveals rules of in vitro evolution. NAR Genom Bioinform 2, lqaa016.

Bai, S., Su, X., Chen, Z., and Han, Z.G. (2025). De Novo Detection of Clonal Structure and Evolution in Single-Cell and Spatial Transcriptomes. Int J Mol Sci 26.

Black, J.R.M., and McGranahan, N. (2021). Genetic and non-genetic clonal diversity in cancer evolution. Nat Rev Cancer 21, 379–392.

Buenrostro, J.D., Wu, B., Litzenburger, U.M., Ruff, D., Gonzales, M.L., Snyder, M.P., Chang, H.Y., and Greenleaf, W.J. (2015). Single-cell chromatin accessibility reveals principles of regulatory variation. Nature 523, 486–490.

Cannoodt, R., Saelens, W., Deconinck, L., and Saeys, Y. (2021). Spearheading future omics analyses using dyngen, a multi-modal simulator of single cells. Nat Commun 12, 3942.

Chen, J., Zhou, Q., Wang, Y., and Ning, K. (2016). Single-cell SNP analyses and interpretations based on RNA-Seq data for colon cancer research. Sci Rep 6, 34420.

Chen, M., Fu, R., Chen, Y., Li, L., and Wang, S.W. (2025). High-resolution, noninvasive single-cell lineage tracing in mice and humans based on DNA methylation epimutations. Nat Methods 22, 488–498.

Funnell, T., O’Flanagan, C.H., Williams, M.J., McPherson, A., McKinney, S., Kabeer, F., Lee, H., Salehi, S., Vazquez-Garcia, I., Shi, H., et al. (2022). Single-cell genomic variation induced by mutational processes in cancer. Nature 612, 106–115.

Gulati, G.S., Sikandar, S.S., Wesche, D.J., Manjunath, A., Bharadwaj, A., Berger, M.J., Ilagan, F., Kuo, A.H., Hsieh, R.W., Cai, S., et al. (2020). Single-cell transcriptional diversity is a hallmark of developmental potential. Science 367, 405–411.

Huang, Y., McCarthy, D.J., and Stegle, O. (2019). Vireo: Bayesian demultiplexing of pooled single-cell RNA-seq data without genotype reference. Genome Biol 20, 273.

Jones, M.G., Khodaverdian, A., Quinn, J.J., Chan, M.M., Hussmann, J.A., Wang, R., Xu, C., Weissman, J.S., and Yosef, N. (2020). Inference of single-cell phylogenies from lineage tracing data using Cassiopeia. Genome Biol 21, 92.

Kharchenko, P.V., Silberstein, L., and Scadden, D.T. (2014). Bayesian approach to single-cell differential expression analysis. Nat Methods 11, 740–742.

La Manno, G., Soldatov, R., Zeisel, A., Braun, E., Hochgerner, H., Petukhov, V., Lidschreiber, K., Kastriti, M.E., Lonnerberg, P., Furlan, A., et al. (2018). RNA velocity of single cells. Nature 560, 494–498.

Li, J., Dang, S.M., Sengupta, S., Schurmann, P., Dost, A.F.M., Moye, A.L., Trovero, M.F., Ahmed, S., Paschini, M., Bhetariya, P.J., et al. (2025a). Organoid modeling reveals the tumorigenic potential of the alveolar progenitor cell state. EMBO J 44, 1804–1828.

Li, Q., Wang, R., Yang, Z., Li, W., Yang, J., Wang, Z., Bai, H., Cui, Y., Tian, Y., Wu, Z., et al. (2022). Molecular profiling of human non-small cell lung cancer by single-cell RNA-seq. Genome Med 14, 87.

Li, S., Wang, K., Wang, X., and Hu, Z. (2025b). Single-cell mitochondrial lineage tracing: Opportunities and challenges. Quantitative Biology 14.

Liu, X., Griffiths, J.I., Bishara, I., Liu, J., Bild, A.H., and Chang, J.T. (2023). Phylogenetic inference from single-cell RNA-seq data. Sci Rep 13, 12854.

Lopez, R., Regier, J., Cole, M.B., Jordan, M.I., and Yosef, N. (2018). Deep generative modeling for single-cell transcriptomics. Nat Methods 15, 1053–1058.

Ludwig, L.S., Lareau, C.A., Ulirsch, J.C., Christian, E., Muus, C., Li, L.H., Pelka, K., Ge, W., Oren, Y., Brack, A., et al. (2019). Lineage Tracing in Humans Enabled by Mitochondrial Mutations and Single-Cell Genomics. Cell 176, 1325–1339 e1322.

Majima, K., Kojima, Y., Minoura, K., Abe, K., Hirose, H., and Shimamura, T. (2024). LineageVAE: reconstructing historical cell states and transcriptomes toward unobserved progenitors. Bioinformatics 40.

McGranahan, N., and Swanton, C. (2017). Clonal Heterogeneity and Tumor Evolution: Past, Present, and the Future. Cell 168, 613–628.

McKenna, A., Findlay, G.M., Gagnon, J.A., Horwitz, M.S., Schier, A.F., and Shendure, J. (2016). Whole-organism lineage tracing by combinatorial and cumulative genome editing. Science 353, aaf7907.

McKenna, A., Hanna, M., Banks, E., Sivachenko, A., Cibulskis, K., Kernytsky, A., Garimella, K., Altshuler, D., Gabriel, S., Daly, M., et al. (2010). The Genome Analysis Toolkit: a MapReduce framework for analyzing next-generation DNA sequencing data. Genome Res 20, 1297–1303.

Muyas, F., Sauer, C.M., Valle-Inclan, J.E., Li, R., Rahbari, R., Mitchell, T.J., Hormoz, S., and Cortes-Ciriano, I. (2024). De novo detection of somatic mutations in high-throughput single-cell profiling data sets. Nat Biotechnol 42, 758–767.

Neftel, C., Laffy, J., Filbin, M.G., Hara, T., Shore, M.E., Rahme, G.J., Richman, A.R., Silverbush, D., Shaw, M.L., Hebert, C.M., et al. (2019). An Integrative Model of Cellular States, Plasticity, and Genetics for Glioblastoma. Cell 178, 835–849 e821.

Nowell, P.C. (1976). The clonal evolution of tumor cell populations. Science 194, 23–28.

Pellegrino, M., Sciambi, A., Treusch, S., Durruthy-Durruthy, R., Gokhale, K., Jacob, J., Chen, T.X., Geis, J.A., Oldham, W., Matthews, J., et al. (2018). High-throughput single-cell DNA sequencing of acute myeloid leukemia tumors with droplet microfluidics. Genome Res 28, 1345–1352.

Petti, A.A., Williams, S.R., Miller, C.A., Fiddes, I.T., Srivatsan, S.N., Chen, D.Y., Fronick, C.C., Fulton, R.S., Church, D.M., and Ley, T.J. (2019). A general approach for detecting expressed mutations in AML cells using single cell RNA-sequencing. Nat Commun 10, 3660.

Qiao, Y., Cheng, T., Miao, Z., Cui, Y., and Tu, J. (2025). Recent Innovations and Technical Advances in High-Throughput Parallel Single-Cell Whole-Genome Sequencing Methods. Small Methods 9, e2400789.

Qin, Z., Yue, M., Tang, S., Wu, F., Sun, H., Li, Y., Zhang, Y., Izumi, H., Huang, H., Wang, W., et al. (2024). EML4-ALK fusions drive lung adeno-to-squamous transition through JAK-STAT activation. J Exp Med 221.

Quinn, J.J., Jones, M.G., Okimoto, R.A., Nanjo, S., Chan, M.M., Yosef, N., Bivona, T.G., and Weissman, J.S. (2021). Single-cell lineages reveal the rates, routes, and drivers of metastasis in cancer xenografts. Science 371.

Raj, B., Gagnon, J.A., and Schier, A.F. (2018). Large-scale reconstruction of cell lineages using single-cell readout of transcriptomes and CRISPR-Cas9 barcodes by scGESTALT. Nat Protoc 13, 2685–2713.

Salvador-Martinez, I., Grillo, M., Averof, M., and Telford, M.J. (2019). Is it possible to reconstruct an accurate cell lineage using CRISPR recorders? Elife 8.

Schiebinger, G., Shu, J., Tabaka, M., Cleary, B., Subramanian, V., Solomon, A., Gould, J., Liu, S., Lin, S., Berube, P., et al. (2019). Optimal-Transport Analysis of Single-Cell Gene Expression Identifies Developmental Trajectories in Reprogramming. Cell 176, 928–943 e922.

Shapiro, E., Biezuner, T., and Linnarsson, S. (2013). Single-cell sequencing-based technologies will revolutionize whole-organism science. Nat Rev Genet 14, 618–630.

Simeonov, K.P., Byrns, C.N., Clark, M.L., Norgard, R.J., Martin, B., Stanger, B.Z., Shendure, J., McKenna, A., and Lengner, C.J. (2021). Single-cell lineage tracing of metastatic cancer reveals selection of hybrid EMT states. Cancer Cell 39, 1150–1162 e1159.

Stahl, P.L., Salmen, F., Vickovic, S., Lundmark, A., Navarro, J.F., Magnusson, J., Giacomello, S., Asp, M., Westholm, J.O., Huss, M., et al. (2016). Visualization and analysis of gene expression in tissue sections by spatial transcriptomics. Science 353, 78–82.

Tang, S., Xue, Y., Qin, Z., Fang, Z., Sun, Y., Yuan, C., Pan, Y., Zhao, Y., Tong, X., Zhang, J., et al. (2023). Counteracting lineage-specific transcription factor network finely tunes lung adeno-to-squamous transdifferentiation through remodeling tumor immune microenvironment. Natl Sci Rev 10, nwad028.

Trapnell, C., Cacchiarelli, D., Grimsby, J., Pokharel, P., Li, S., Morse, M., Lennon, N.J., Livak, K.J., Mikkelsen, T.S., and Rinn, J.L. (2014). The dynamics and regulators of cell fate decisions are revealed by pseudotemporal ordering of single cells. Nat Biotechnol 32, 381–386.

Valecha, M., and Posada, D. (2022). Somatic variant calling from single-cell DNA sequencing data. Comput Struct Biotechnol J 20, 2978–2985.

Vogelstein, B., Papadopoulos, N., Velculescu, V.E., Zhou, S., Diaz, L.A., Jr., and Kinzler, K.W. (2013). Cancer genome landscapes. Science 339, 1546–1558.

Vu, T.N., Nguyen, H.N., Calza, S., Kalari, K.R., Wang, L., and Pawitan, Y. (2019). Cell-level somatic mutation detection from single-cell RNA sequencing. Bioinformatics 35, 4679–4687.

Wagner, D.E., and Klein, A.M. (2020). Lineage tracing meets single-cell omics: opportunities and challenges. Nat Rev Genet 21, 410–427.

Wang, Z., Li, Z., Zhou, K., Wang, C., Jiang, L., Zhang, L., Yang, Y., Luo, W., Qiao, W., Wang, G., et al. (2021). Deciphering cell lineage specification of human lung adenocarcinoma with single-cell RNA sequencing. Nat Commun 12, 6500.

Wang, Z., Zhu, G., Tang, P., Wang, Y., Luo, W., Song, W., Pan, Z., Zheng, B., Jiang, Y., Xiao, D., et al. (2025). SLPI(+) AT2-Like Cells Orchestrate Lung Adenocarcinoma Invasion via Wnt Pathway Activation and Stromal Crosstalk in a Spatially Defined Margin Niche. Adv Sci (Weinh), e16580.

Weber, L.L., Zhang, C., Ochoa, I., and El-Kebir, M. (2023). Phertilizer: Growing a clonal tree from ultra-low coverage single-cell DNA sequencing of tumors. PLoS Comput Biol 19, e1011544.

Wei, G., Zhang, X., Liu, S., Hou, W., and Dai, Z. (2024). Comprehensive data mining reveals RTK/RAS signaling pathway as a promoter of prostate cancer lineage plasticity through transcription factors and CNV. Sci Rep 14, 11688.

Weinreb, C., Wolock, S., Tusi, B.K., Socolovsky, M., and Klein, A.M. (2018). Fundamental limits on dynamic inference from single-cell snapshots. Proc Natl Acad Sci U S A 115, E2467–E2476.

Wiens, M., Farahani, H., Scott, R.W., Underhill, T.M., and Bashashati, A. (2024). Benchmarking bulk and single-cell variant-calling approaches on Chromium scRNA-seq and scATAC-seq libraries. Genome Res 34, 1196–1210.

Xu, J., Nuno, K., Litzenburger, U.M., Qi, Y., Corces, M.R., Majeti, R., and Chang, H.Y. (2019). Single-cell lineage tracing by endogenous mutations enriched in transposase accessible mitochondrial DNA. Elife 8.

Yang, D., Jones, M.G., Naranjo, S., Rideout, W.M., 3rd, Min, K.H.J., Ho, R., Wu, W., Replogle, J.M., Page, J.L., Quinn, J.J., et al. (2022). Lineage tracing reveals the phylodynamics, plasticity, and paths of tumor evolution. Cell 185, 1905–1923 e1925.

Zhang, T., Jia, H., Song, T., Lv, L., Gulhan, D.C., Wang, H., Guo, W., Xi, R., Guo, H., and Shen, N. (2023). De novo identification of expressed cancer somatic mutations from single-cell RNA sequencing data. Genome Med 15, 115.

