## Supplementary Tables and Figures VAETracer for "VAETracer: Mutation-Guided Lineage Reconstruction and Generational State Inference from scRNA-seq"

**Supplementary Table 1: Correlation between predicted and true values under simulated simple and lineage data.**

Spearman correlation coefficients (rho) were computed between inferred and ground-truth cellular mutation profiles (CMM) across different method's simple combinations, data types, and parameter settings. Significance levels are indicated by asterisks: * $p < 0.05$, ** $p < 0.01$, *** $p < 0.001$.

- data_type: Type of simulated data (simple or lineage-structured).
- pipe_type: Method combination used for CMM inference; VAE_rev indicates that the input to VAE was transposed.
- value_type: Target variable (N or P).
- method_type: Component model whose output is evaluated.
- Beta: Beta distribution parameters used to generate the ground-truth p values.

Note: Simulated data contain 10% mask (missing values). Each condition has only one replicate.

Results show that models perform better under lower mutation rates, where more informative signals are preserved. In contrast, when all sites have high mutation probabilities, the input contains less effective information, making it harder for models to capture meaningful patterns.

| **type** | | | | **beta** | | | |
|---|---|---|---|---|---|---|---|
| **data_type** | pipe_type | value_type | method_type | Beta(1,32) | Beta(0.1,0.5) | Beta(8,4) | Beta1(2,38)+ Beta2(16,4) |
| **simple** | gNMF + FT | N | gNMF | 0.997*** | 0.989*** | 0.410*** | 0.871*** |
| | | | FT | 0.997*** | 0.989*** | 0.409*** | 0.872*** |
| | | P | gNMF | 0.973*** | 0.882*** | 0.837*** | 0.893*** |
| | | | FT | 0.997*** | 0.980*** | 0.906*** | 0.987*** |
| | VAE + FT | N | VAE | 0.981*** | 0.846*** | 0.358*** | 0.872*** |
| | | | FT | 0.981*** | 0.846*** | 0.360*** | 0.872*** |
| | | P | VAE | 0.840*** | 0.723*** | 0.836*** | 0.924*** |
| | | | FT | 0.993*** | 0.774*** | 0.904*** | 0.990*** |
| | VAE_rev + FT | N | VAE | 0.990*** | 0.984*** | 0.532*** | 0.903*** |
| | | | FT | 0.990*** | 0.983*** | 0.529*** | 0.902*** |
| | | P | VAE | 0.927*** | 0.823*** | 0.041 | 0.863*** |
| | | | FT | 0.995*** | 0.957*** | 0.041 | 0.859*** |
| **lineage** | gNMF + FT | N | gNMF | 0.915*** | 0.638*** | -0.021 | 0.884*** |
| | | | FT | 0.914*** | 0.637*** | 0.000 | 0.882*** |
| | | P | gNMF | 0.793*** | 0.679*** | 0.004 | 0.749*** |
| | | | FT | 0.755*** | 0.684*** | 0.004 | 0.758*** |

| | | | | | | | |
|---|---|---|---|---|---|---|---|
| | VAE + FT | N | VAE | 0.877*** | 0.426*** | -0.042 | 0.851*** |
| | | | FT | 0.877*** | 0.427*** | 0.000 | 0.851*** |
| | | P | VAE | 0.794*** | 0.679*** | 0.002 | 0.783*** |
| | | | FT | 0.755*** | 0.681*** | 0.002 | 0.800*** |
| | VAE_rev + FT | N | VAE | 0.906*** | 0.883*** | -0.028 | 0.820*** |
| | | | FT | 0.905*** | 0.883*** | 0.000 | 0.813*** |
| | | P | VAE | 0.782*** | 0.725*** | 0.037 | 0.518*** |
| | | | FT | 0.748*** | 0.744*** | 0.037 | 0.485*** |

**Supplementary Table 2: Performance comparison of complex method combinations under three representative beta distributions with repeated simulations.**

Spearman correlation coefficients (rho) are reported as mean +- SEM across 50 independent replicates, focusing on three beta settings with relatively low mutation rates (excluding high-mutation-only conditions). Complex pipelines were tested, including using gNMF output as input or reference for VAE training.

- data_type: Type of simulated data (simple or lineage-structured).
- pipe_type: Complex method combination.
- value_type: Target variable (N or P).
- method_type: Evaluated component model.
- VAE_type: Whether to transpose input for VAE
- Beta: Beta distribution parameters used to generate the ground-truth p values.

Key findings:

1. gNMF achieves better overall performance compared to VAE-based approaches.
2. VAE performs well on low-mutation events, but convergence becomes unstable when reference inputs are provided, often leading to suboptimal results.
3. VAE_rev shows superior performance, suggesting that cell-dimensional structure is more complex, and the inferred N from the latent representations is more sensitive than P to input patterns and noise.

| type | | | | | beta | | |
|---|---|---|---|---|---|---|---|
| **data type** | **pipe_type** | **value type** | **method type** | **VAE_type** | **Beta(1,32)** | **Beta(0.1,0.5)** | **Beta1(2,38) +Beta2(16,4)** |
| **simple** | gNMF+FT | N | gNMF | | 0.99161 ± 1.1e-04 | 0.95959 ± 5.8e-04 | 0.87106 ± 6.0e-04 |
| | | P | | | 0.998486 ± 1.0e-05 | 0.94481 ± 7.5e-04 | 0.96784 ± 5.6e-04 |
| | VAE+FT | N | VAE | VAE | 0.95921 ± 5.0e-04 | 0.86329 ± 0.00520 | 0.91786 ± 0.00326 |
| | | | | VAE_rev | 0.96563 ± 4.8e-04 | 0.95876 ± 0.00263 | 0.88097 ± 0.00318 |
| | | P | | VAE | 0.998474 ± 1.1e-05 | 0.94478 ± 7.6e-04 | 0.96505 ± 6.4e-04 |
| | | | | VAE_rev | 0.99393 ± 2.0e-04 | 0.88441 ± 0.00160 | 0.87356 ± 8.2e-04 |
| | gNMF+ VAE+FT | N | | VAE | 0.5446 ± 0.0591 | 0.2927 ± 0.0315 | 0.7268 ± 0.0155 |
| | | | | VAE_rev | 0.98313 ± 1.5e-04 | 0.96814 ± 5.0e-04 | 0.89599 ± 4.9e-04 |
| | | P | | VAE | 0.998484 ± 1.0e-05 | 0.94481 ± 7.5e-04 | 0.96784 ± 5.6e-04 |
| | | | | VAE_rev | 0.99304 ± 2.2e-04 | 0.87877 ± 0.00196 | 0.88353 ± 9.9e-04 |
| | gNMF+FT+VAE | N | | VAE | 0.84658 ± 0.00373 | 0.91965 ± 0.00247 | 0.22758 ± 0.00833 |
| | | | | VAE_rev | 0.99132 ± 1.1e-04 | 0.95883 ± 5.7e-04 | 0.87041 ± 6.2e-04 |
| | | P | | VAE | 0.998444 ± 1.1e-05 | 0.94481 ± 7.5e-04 | 0.97133 ± 4.5e-04 |
| | | | | VAE_rev | 0.8998 ± 0.0173 | 0.78566 ± 0.00953 | 0.88147 ± 0.00133 |
| **lineage** | gNMF+FT | N | gNMF | | 0.95606 ± 0.00187 | 0.87158 ± 0.00489 | 0.89089 ± 0.00292 |
| | | P | | | 0.8814 ± 0.0129 | 0.92468 ± 0.00613 | 0.80562 ± 0.00675 |
| | VAE+FT | N | VAE | VAE | 0.8392 ± 0.0211 | 0.2107 ± 0.0214 | 0.3621 ± 0.0362 |
| | | | | VAE_rev | 0.92490 ± 0.00345 | 0.86232 ± 0.00849 | 0.85188 ± 0.00358 |
| | | P | | VAE | 0.8783 ± 0.0147 | 0.92469 ± 0.00604 | 0.79748 ± 0.00723 |

| | | | | | | | |
|---|---|---|---|---|---|---|---|
| | | | | VAE_rev | 0.8714 ± 0.0160 | 0.83558 ± 0.00531 | 0.6163 ± 0.0124 |
| | gNMF+ VAE+FT | N | | VAE | 0.7346 ± 0.0228 | 0.0741 ± 0.0262 | 0.3483 ± 0.0348 |
| | | | | VAE_rev | 0.95000 ± 0.00197 | 0.87022 ± 0.00520 | 0.90274 ± 0.00272 |
| | | P | | VAE | 0.8814 ± 0.0129 | 0.92467 ± 0.00613 | 0.80562 ± 0.00675 |
| | | | | VAE_rev | 0.8720 ± 0.0162 | 0.83574 ± 0.00495 | 0.7257 ± 0.0103 |
| | gNMF+FT+VAE | N | | VAE | 0.6646 ± 0.0242 | 0.5917 ± 0.0169 | 0.4320 ± 0.0291 |
| | | | | VAE_rev | 0.95456 ± 0.00187 | 0.86382 ± 0.00514 | 0.88209 ± 0.00296 |
| | | P | | VAE | 0.8741 ± 0.0126 | 0.92463 ± 0.00613 | 0.80257 ± 0.00713 |
| | | | | VAE_rev | 0.7437 ± 0.0256 | 0.7467 ± 0.0277 | 0.74734 ± 0.00799 |

**Supplementary Table 3: Performance of different method combinations on lineage-simulated data over time under Beta(1,32).**

Results are based on temporally segmented lineage-structured data simulated with Beta(1,32), where mutation rates follow a low-mean distribution. Various method combinations were applied to infer cellular mutation profiles (CMM) at different time stages. The table summarizes the Spearman correlation coefficients (rho) between predicted and true values, showing how model performance evolves along the simulated lineage trajectory.

| pipe_type | value_type | method_type | VAE_type | Correlation |
|---|---|---|---|---|
| gNMF+FT | N | gNMF | | 0.996*** |
| | P | | | 0.965*** |
| | N | FT | | 0.995*** |
| | P | | | 0.965*** |
| VAE+FT | N | VAE | VAE | 0.990*** |
| | | FT | | 0.990*** |
| | P | VAE | | 0.965*** |
| | | FT | | 0.965*** |
| | N | VAE | VAE_rev | 0.994*** |
| | | FT | | 0.994*** |
| | P | VAE | | 0.963*** |
| | | FT | | 0.964*** |
| gNMF+VAE+FT | N | VAE | VAE | 0.984*** |
| | | FT | | 0.984*** |
| | P | VAE | | 0.965*** |
| | | FT | | 0.965*** |
| | N | VAE | VAE_rev | 0.991*** |
| | | FT | | 0.991*** |
| | P | VAE | | 0.963*** |
| | | FT | | 0.964*** |

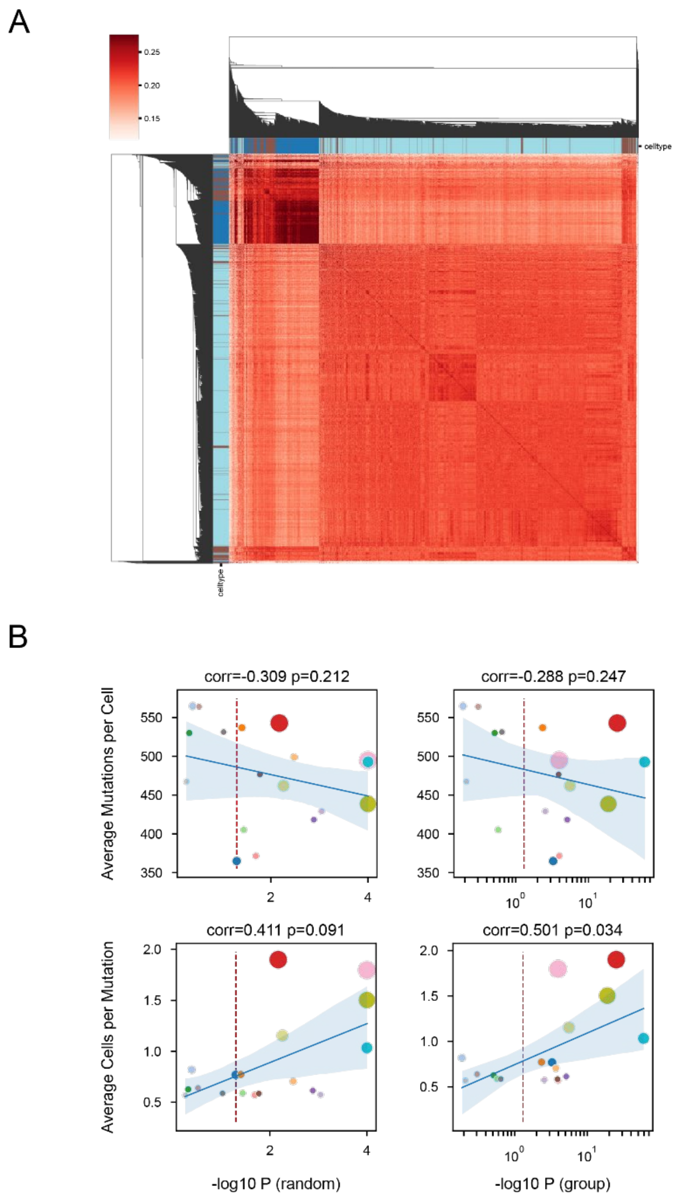


**Supplementary Figure 1:**

Supplementary Figure 1: scRNA-seq-inferred mutations reflect cellular lineage structure.

(A) Heatmap of pairwise correlations between mutation profiles across single cells from the LUAS dataset. Cells are ordered by hierarchical clustering; dendrograms reflect similarity in mutation patterns. Color bar indicates correlation strength. Cell types are labeled along the top: adenocarcinoma (ADC, dark blue), squamous cell carcinoma (SCC, light blue), mixed (Mix, brown).

(B) Clone-level phylogenetic significance (-log10 P) plotted against two mutation quality metrics. Top panels show no significant correlation between average mutations per cell and significance under either random (left) or group-based (right) null models. Bottom panels reveal a positive trend between average cells per mutation and significance, indicating that mutations shared across more cells are more likely to capture true lineage relationships. Each point represents one clone. Red dashed lines denote P = 0.05 threshold.

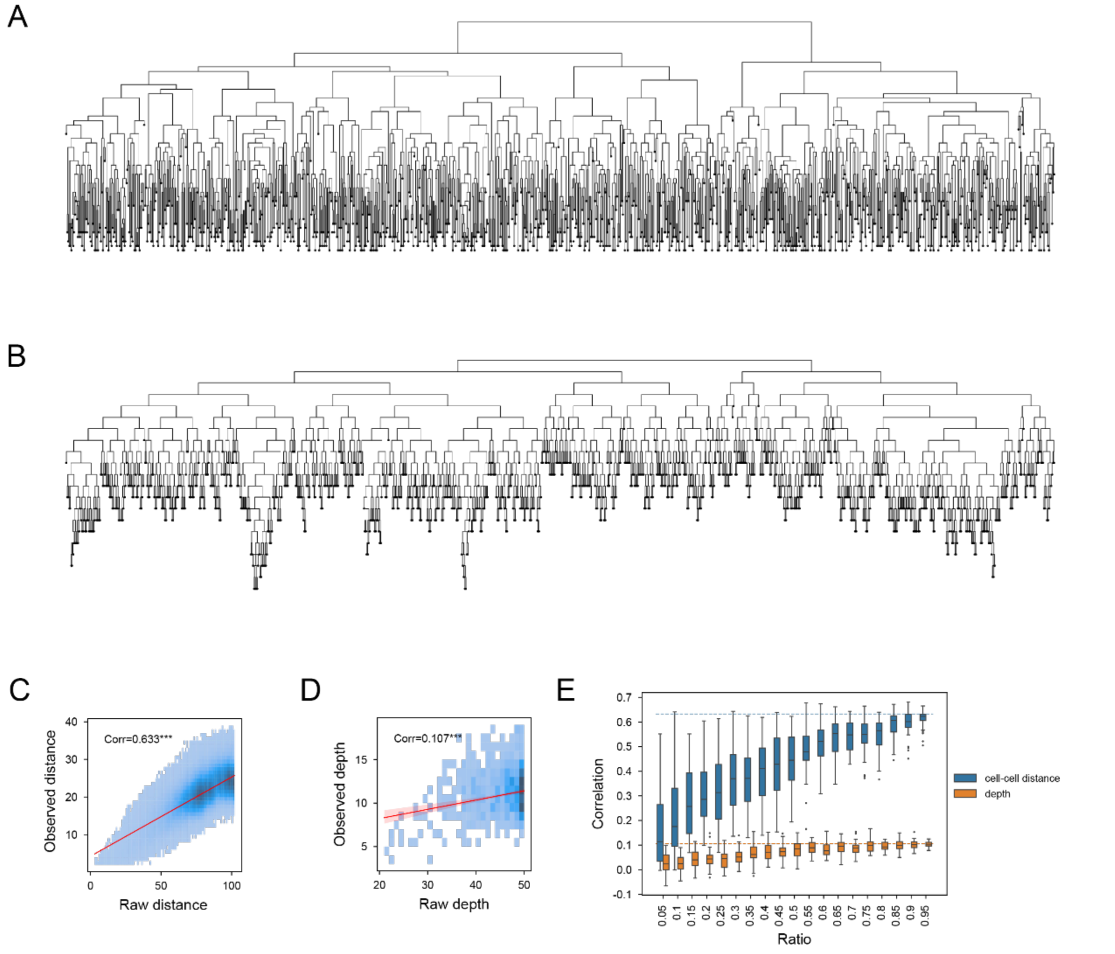


**Supplementary Figure 2: Traditional tree-based lineage inference is sensitive to sampling completeness and fails to robustly recover developmental progression despite capturing cell-cell relationships.**

(A) Simulated true lineage tree with 1000 terminal cells (leaf nodes) and internal ancestral nodes, reflecting a branching developmental process. (B) Simplified tree derived from (A) by setting all edge lengths to 1, which represents the assumption of uniform mutation accumulation rate in traditional phylogenetic methods. This structure serves as the ground truth for evaluating distance and depth estimation. (C,D) Comparison between observed (inferred) and raw (true) cell-cell distances (C) and cell depths (D) under full sampling. While cell-cell distances show moderate correlation with truth, depth estimates exhibit very weak agreement, indicating that traditional tree reconstruction poorly recovers developmental progression even when pairwise relationships are preserved. (E) Correlation between inferred and true metrics across varying ancestral node sampling ratios (x-axis). Both cell-cell distance (blue boxes) and depth (orange boxes) decrease with lower sampling rates, but depth correlation remains consistently near zero across all conditions, while distance correlation declines more gradually. This highlights the fundamental fragility of depth estimation under sparse sampling as a key limitation of tree-based approaches. *Correlations are computed using Spearman's rank correlation; significance levels: *<0.05, **<0.01, ***<0.001.*

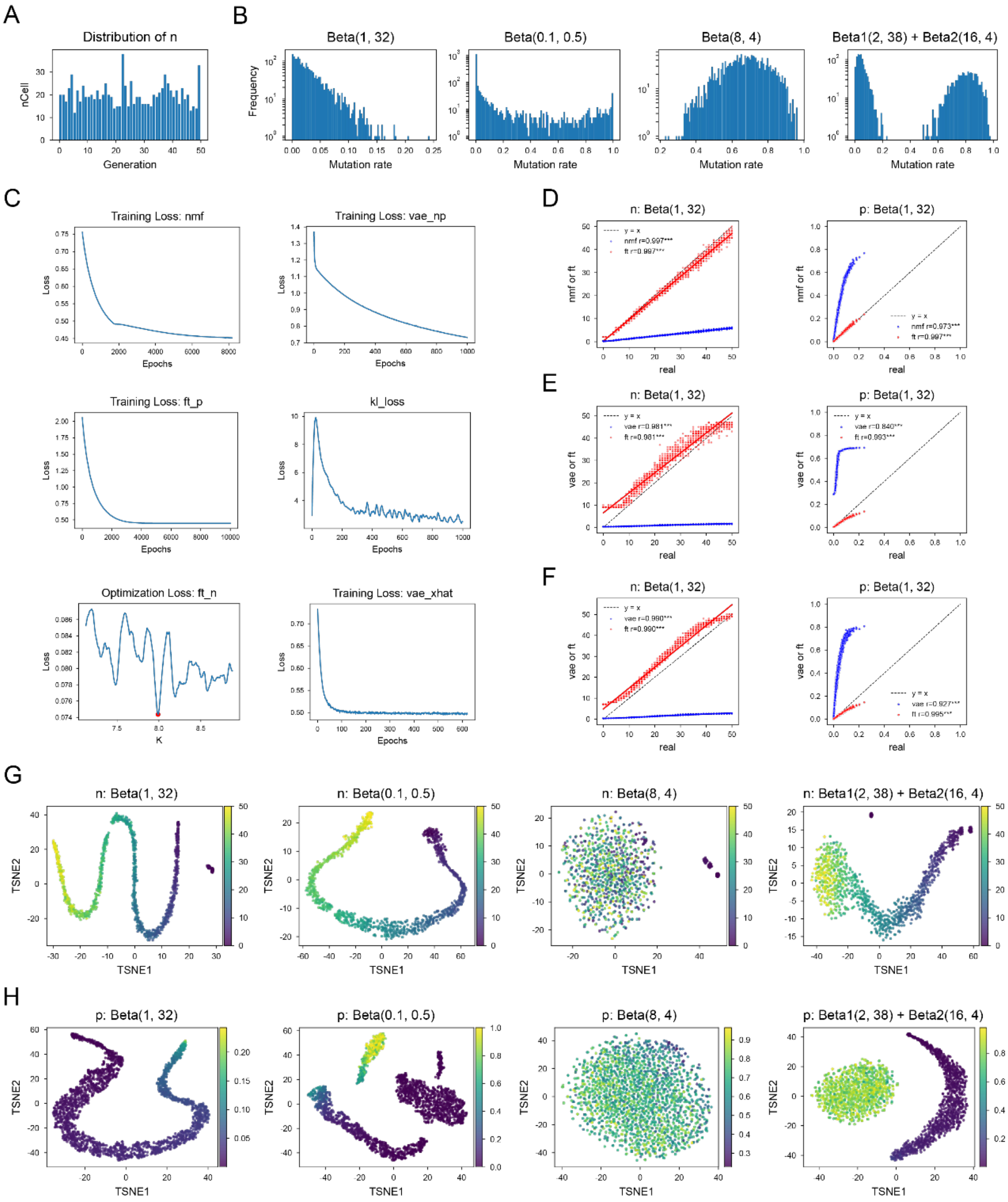


**Supplementary Figure 3:** scMut accurately recovers CGI (N) and site-specific mutation rates (P) from sparse, noisy mutation matrices under diverse simulation settings.

(A) Distribution of true N across 1000 simulated cells.

(B) Four distinct parameterizations of P for 2000 sites, modeled using Beta distributions: Beta(1,32), Beta(0.1,0.5), Beta(8,4), and a bimodal mixture Beta(2,38)+Beta(16,4).

(C) Training loss curves for each component of scMut: gNMF (nmf), VAE (mode-np, vae_np), KL divergence (kl_loss), fine-tuning for N and P (ft_n / ft_p), and VAE mode-xhat loss (vae_xhat).

(D-F) Comparison between inferred and true N and P values under the Beta(1,32) setting, using different module combinations: (D) gNMF with fine-tuning (FT), (E)

VAE with FT, and (F) VAE_rev with FT. N is recovered with high linear correlation to truth; FT refines estimates to near-perfect agreement. For P, relative ordering is preserved but nonlinear scaling is observed; FT corrects this bias and accurately recovers low-mutation-rate sites.
(G) t-SNE visualization of the latent representation $Z_m$ learned by VAE, colored by true N values. Each panel corresponds to one P distribution in (B). The embedding reveals continuous developmental trajectories aligned with increasing N.
(H) t-SNE visualization of $Z_m$ colored by true P values. The latent space preserves the global structure of mutation rate heterogeneity across sites. Note: These embeddings are derived from VAE_rev, a transposed input variant of the VAE used to decode site-level patterns from cell-level representations.
*For N, correlations were assessed using linear regression (scipy.stats.linregress); for P, Spearman's rank correlation was used. Significance levels: *<0.05, **<0.01, ***<0.001.*

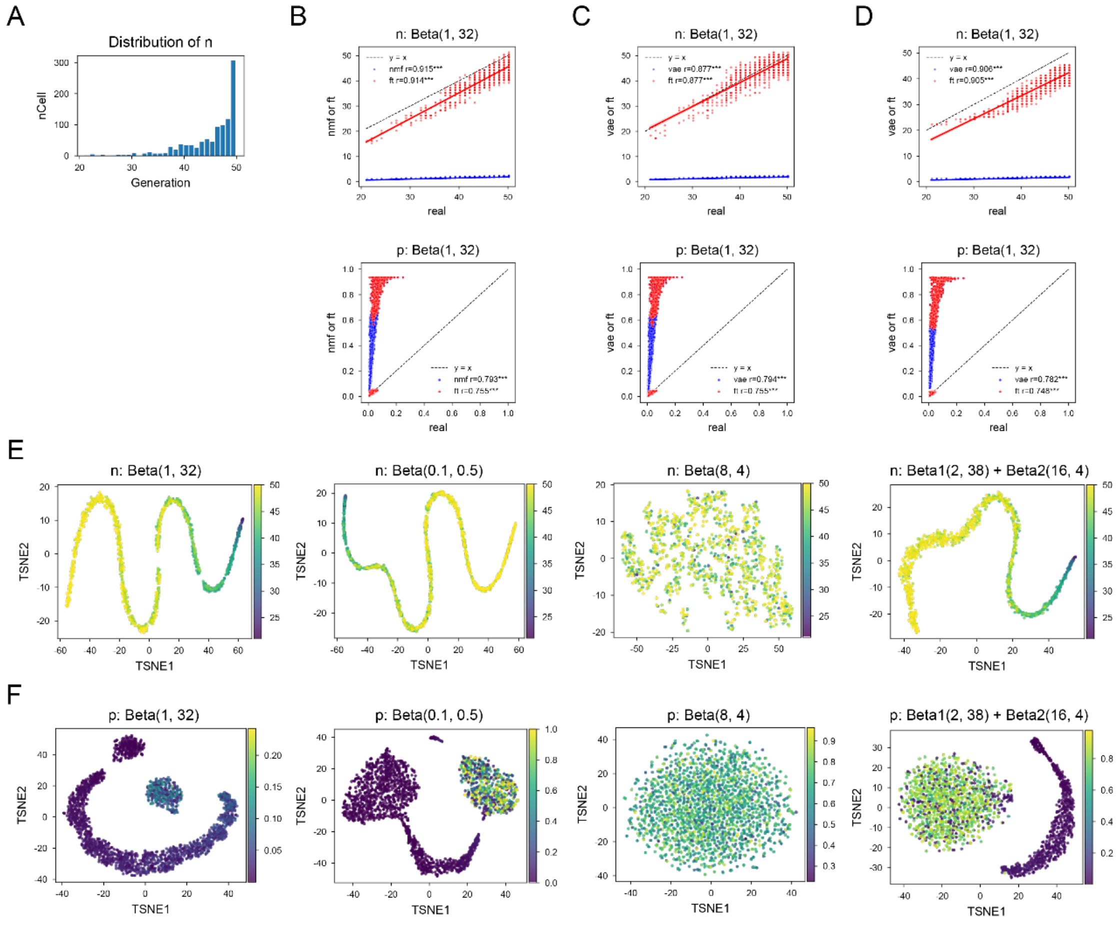


**Supplementary Figure4: scMut robustly recovers CGI (N) and site-specific mutation rates (P) from phylogenetically simulated mutation matrices under biologically realistic conditions.**

(A) Distribution of true N across 1000 cells sampled from a phylogenetic tree with maximum depth of 50 generations.

(B-D) Comparison between inferred and true N and P values under the Beta(1,32) setting, using different module combinations.

(E,F) t-SNE visualization of the latent representation $Z_m$ learned by VAE (E) or VAE_rev (F), colored by true N (E) or P (F) values. The embedding reveals continuous developmental trajectories, except for Beta(8,4).

*(B-F) Methods as in Supplementary Figure 3.*

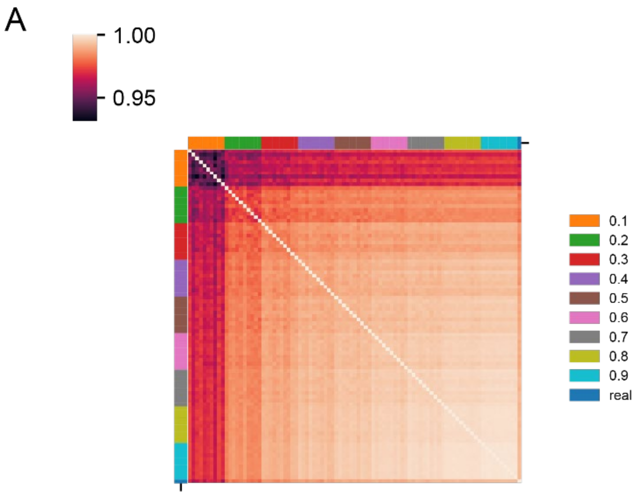


**Supplementary Figure 5: scMut maintains high concordance in CGI estimates across different levels of data sparsity.**

(A) Heatmap showing pairwise Spearman correlation between inferred N values under different missingness levels (0.1 to 0.9) and the true N. Correlation values range from 0.95 to 1.00, indicating strong stability of N estimates even at 90% missing data.

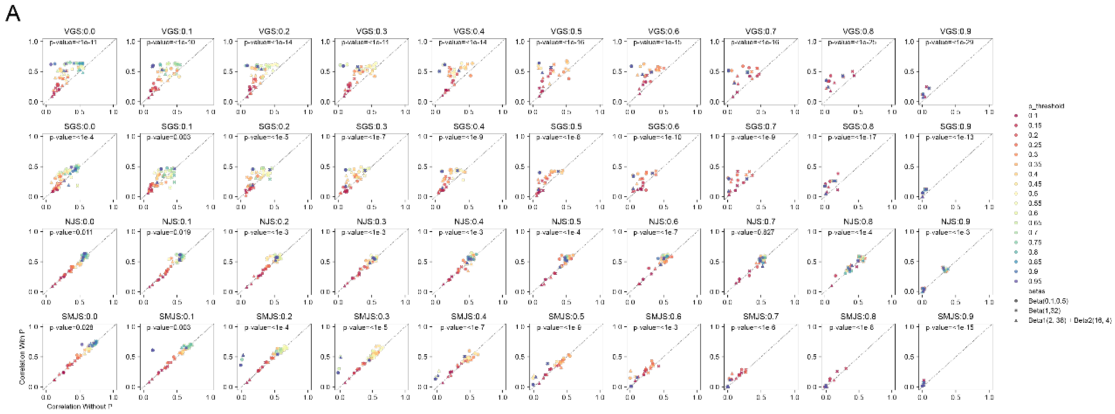

**Supplementary Figure 6: Inferred site-specific mutation rates (P) improve the accuracy of conventional phylogenetic tree reconstruction under heterogeneous mutation landscapes.**

(A) Scatter plots comparing cell-cell distance correlation between true and inferred trees, with (y-axis) and without (x-axis) using scMut-derived P as site weights. Each row corresponds to a different tree-building algorithm implemented in Cassiopeia: VanillaGreedySolver (VGS), SpectralGreedySolver (SGS), NeighborJoiningSolver (NJS), and SharedMutationJoiningSolver (SMJS). Columns represent increasing levels of data sparsity (0.0 to 0.9), where values indicate the proportion of masked entries in the input mutation matrix. Points are colored by P-threshold (legend on right), indicating sites included in weighted distance calculation; point shapes correspond to three distinct P generation distributions.

*Correlations were computed using Pearson's correlation to align with Mantel test assumptions. Statistical significance of improvement (y > x) was assessed via one-sample t-test (scipy.stats.ttest_1samp) on y-x differences.*

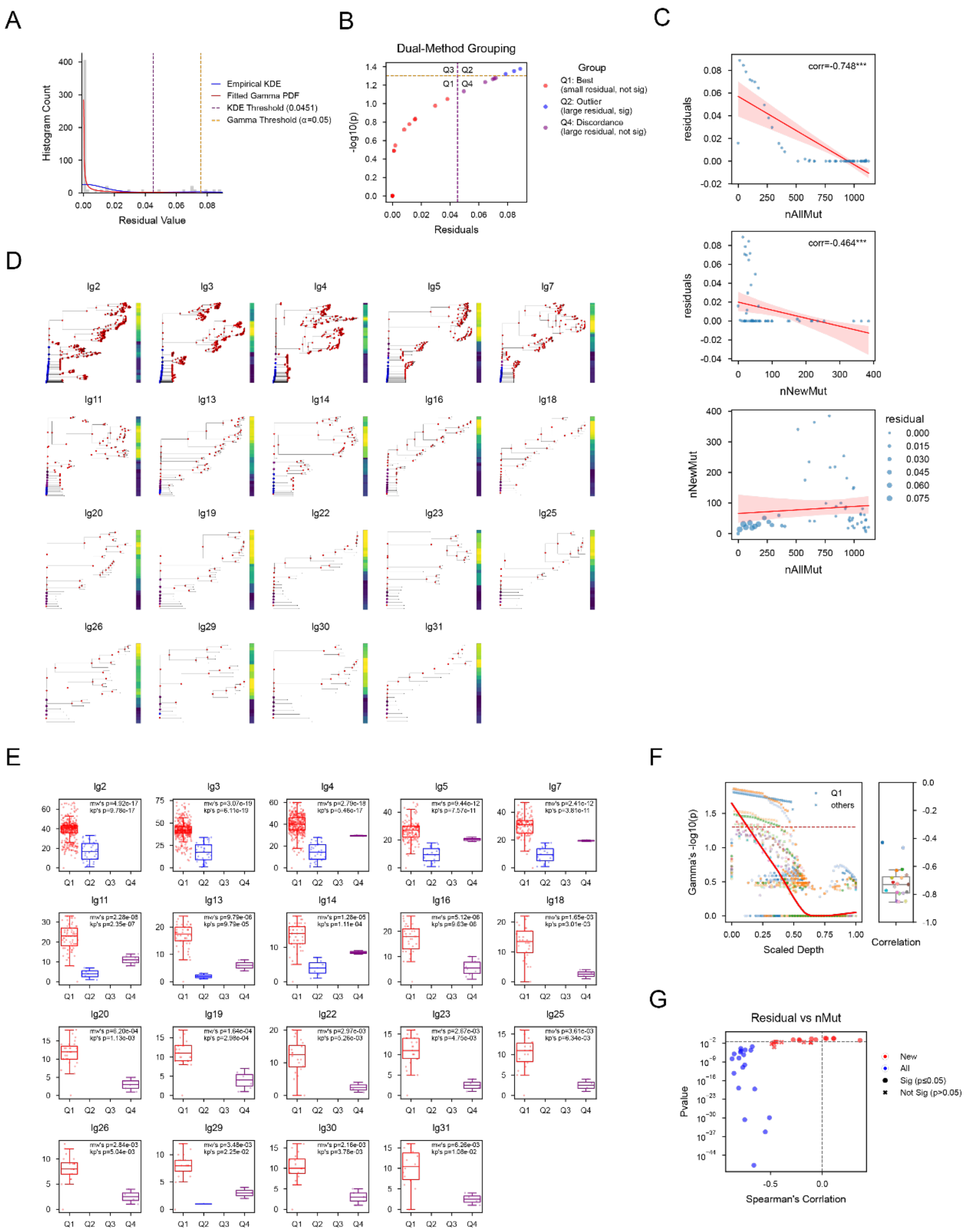

**Supplementary Figure 7: scMut enables time-resolved lineage tree reconstruction by integrating inferred CGI (N) with phylogenetic topology, revealing node-level confidence and biological correlates of estimation stability.** To enable quantitative lineage analysis, we combine traditional tree reconstruction (Cassiopeia) with scMut: ancestral nodes are imputed via phylogeny-aware methods, then all nodes (ancestral + terminal) are input into scMut to infer CGI (N) and latent representation $Z_m$. Edge lengths are computed as $\Delta N = N\_child - N\_parent$; negative $\Delta N$ values are constrained to a minimum of 0.001. Residuals were defined as the difference between the original N estimate from scMut (N_raw) and the updated N

value after enforcing non-negative edge lengths (N_now). Nodes with large residuals are considered unstable or unreliable.
(A) Distribution of residual values across ancestral nodes in Clone-13. Two independent methods identify outlier nodes:
1.Kernel density estimation (KDE) fits a bimodal distribution; the purple vertical line marks the threshold separating high-confidence (small residual) and low-confidence (large residual) nodes.
2.Gamma distribution fitting estimates significance of deviation; the orange horizontal line indicates $p = 0.05$ threshold.
(B) Dual-method grouping of ancestral nodes based on residual magnitude and significance:
Q1: Small residual, not significant (high confidence)
Q2: Large residual, significant (low confidence)
Q3: Discordant classification (small residual, significant, not observed in data)
Q4: Discordant classification (large residual, not significant)
(C) Scatter plots for Clone-13 showing:
Top: Negative correlation between residual and total mutations per node.
Middle: Weaker negative correlation between residual and new mutations acquired from parent.
Bottom: Point distributions of total vs. new mutations per node.
(D) Time-resolved lineage trees for all 19 clones, with edge lengths proportional to $\Delta N$. Node colors indicate group membership (Q1-Q4), highlighting regions of high (Q1) and low (Q2/Q4) estimation confidence.
(E) Boxplots of node's depth for each group (Q1-Q4) across all clones. Q1 nodes show deeper positions, consistent with stable estimation at later developmental stages. For each clone, statistical tests are shown:
Kruskal-Wallis test (kp) compares depth distributions across Q1-Q4 groups.
Mann-Whitney U test (mw) compares Q1 vs. non-Q1 nodes.
(F) Relationship between scaled depth and gamma-distributed p-value for residual across all clones. Nodes deeper in the tree tend to have higher p-values (less significant deviation), supporting greater stability at later branches.
(G) Global validation of (C): Correlation between residual and total/new mutations per node across all clones. Strong negative correlation confirms that estimation reliability increases with mutation burden.
*Spearman's rank correlation was used. Significance levels: *<0.05, **<0.01, ***<0.001.*

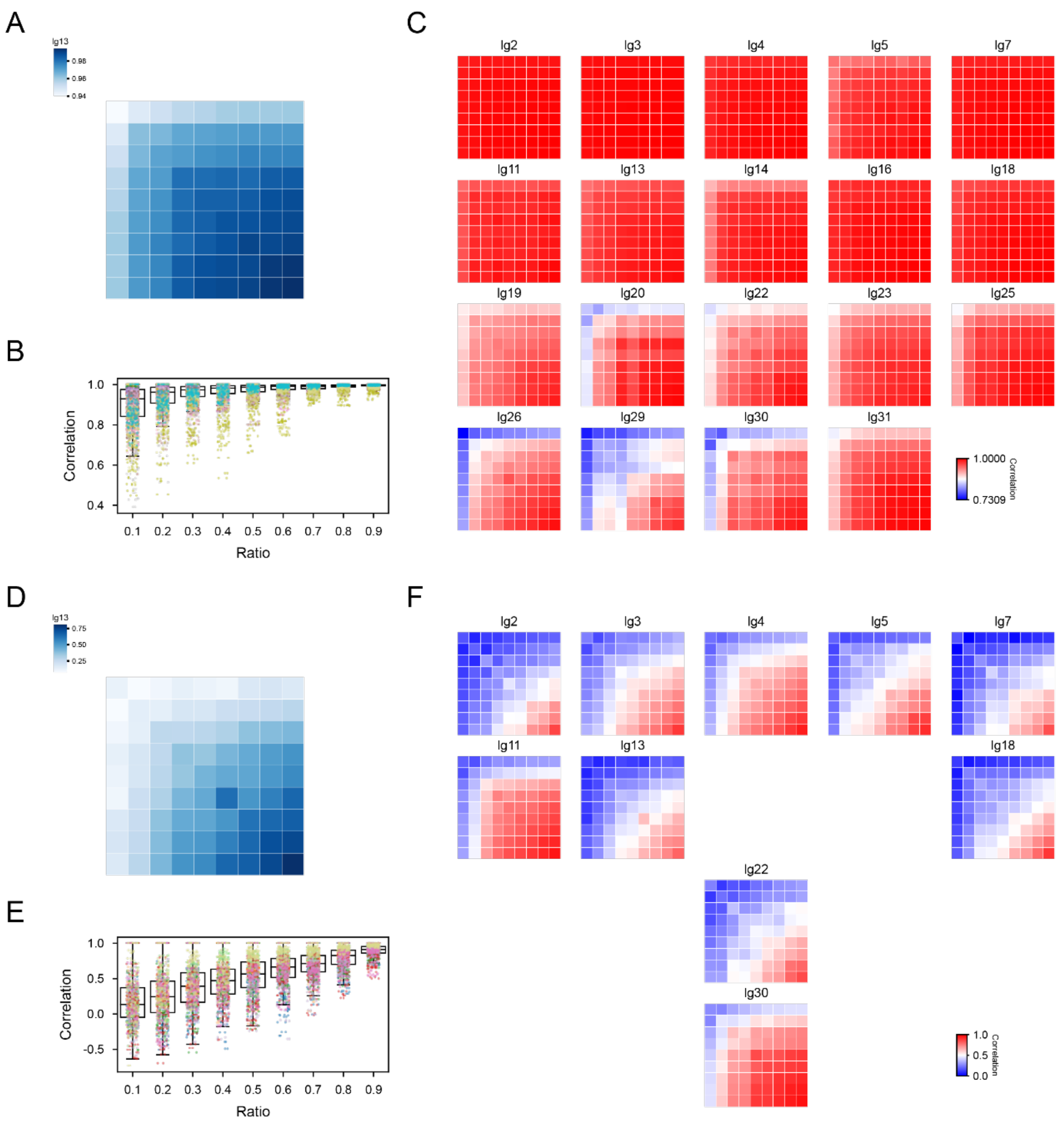


**Supplementary Figure 8: scMut produces highly reproducible CGI (N) estimates for mutation profiles from scRNA-seq under downsampling and generalizes to Cas9-based lineage recording data.**

(A) Heatmap of mean pairwise correlation between inferred N values across different downsampling ratios (0.1 to 0.9, step 0.1) for Clone-13. High correlation (blue) indicates strong consistency in N estimation despite data sparsity.

(B) Boxplots showing distribution of pairwise correlations within each downsampling ratio across all 19 clones. Correlations remain high (>0.8) even at low sampling rates, demonstrating robustness to missing data.

(C) Heatmaps of mean pairwise correlation across all clones. Most clones show consistently high correlation (red), with a few exhibiting moderate variability (lighter shades), reflecting clone-specific noise levels.

(D) Heatmap of mean pairwise correlation between inferred N values across downsampling ratios for Cas9-based editing data in Clone-13. Similar to (A), high correlation indicates stable N inference under downsampling.

(E) Boxplots showing distribution of pairwise correlations within each downsampling ratio across all Cas9 clones. Correlations are generally lower than in scRNA-seq data but still show clear positive trends, indicating qualitative consistency.
(F) Heatmaps of mean pairwise correlation across all Cas9 clones. While absolute correlation values vary more than in scRNA-seq data, the overall pattern of stability under downsampling is preserved.
*Correlations computed using Spearman's rank correlation; significance levels: *<0.05, **<0.01, ***<0.001.*

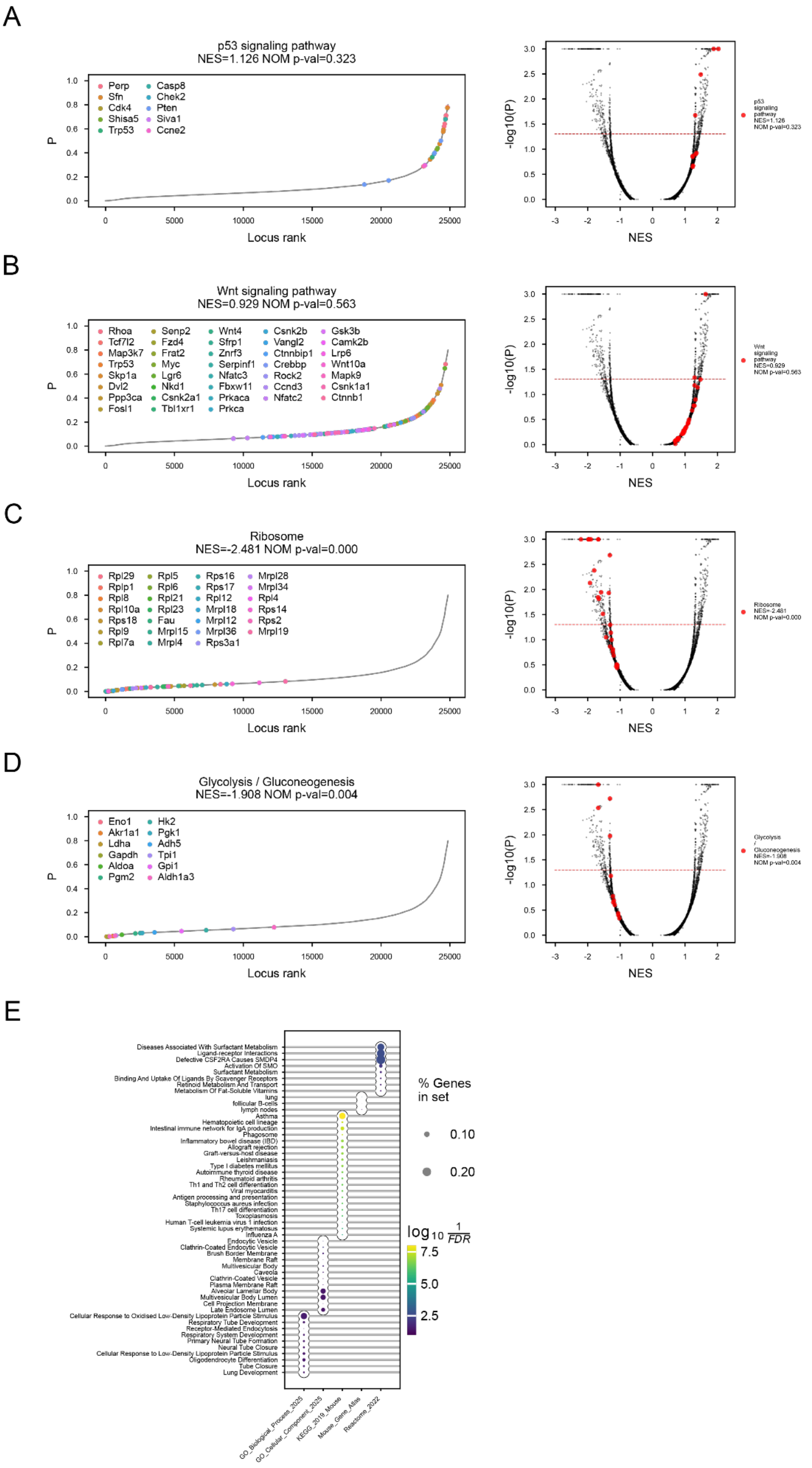


**Supplementary Figure 9: Functional annotation of site-specific mutation rates (P) reveals lineage-associated molecular programs in the LUAS dataset.**

(A-D) Gene set enrichment analysis of inferred P values for four representative pathways using GSEA (gseapy.prerank), ordered by locus rank (left) and visualized via enrichment plots (right).
(A,B) Pathways enriched for high mutation burden: p53 signaling and Wnt signaling, consistent with known roles in tumor proliferation and genomic instability.
(C,D) Pathways enriched for low mutation burden (conserved regions): Ribosome and Glycolysis / Gluconeogenesis, indicating that core metabolic functions remain highly conserved in tumor evolution.
(E) Enrichment analysis of cell-type-specific sites using Enrichr (gseapy.enrichr). Pathways are ranked by -log10(FDR), with dot size indicating percentage of genes in the set. Top enriched terms are shown, aligning with known LUAS biology and supporting scMut's ability to uncover mutation-centric functional programs.

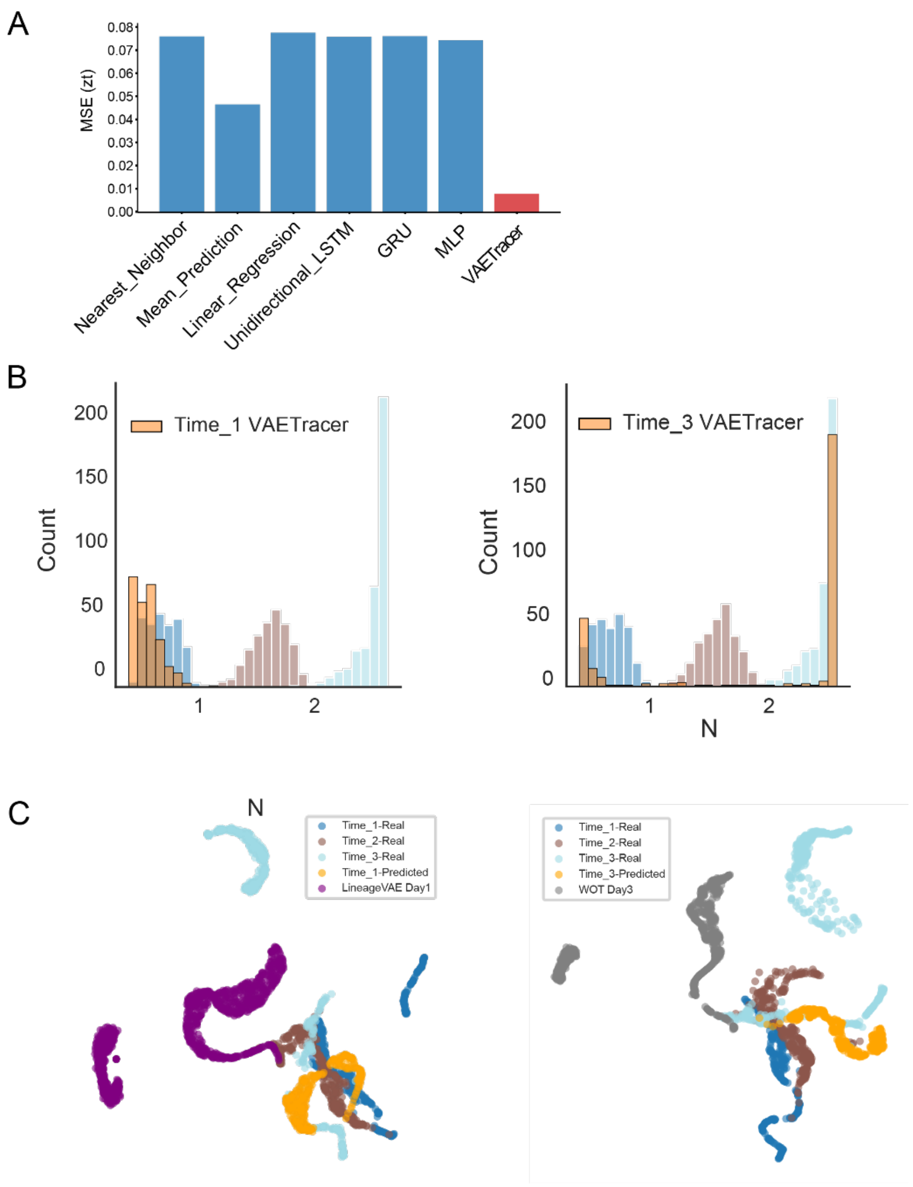


**Supplementary Figure 10: Validation of mutation-informed temporal inference and latent-state prediction.**

(A) Benchmark evaluation of latent-state prediction accuracy on simulated datasets. Multiple baseline methods were compared for predicting missing temporal states, including nearest-neighbor, mean prediction, linear interpolation, linear regression, multilayer perceptron (MLP), unidirectional LSTM, and GRU models. Baseline models were trained to predict latent representations using the concatenation of mutational and transcriptional latent representations ($Z_{mt} + Z_{xt}$), depending on the method. Prediction performance was quantified using mean squared error (MSE) between predicted and ground-truth latent representations at the target time point, demonstrating the advantage of joint mutation-transcription modeling for accurate temporal inference.

(B) Validation of CGI (N) prediction on simulated data under forward and backward inference settings. Left: Forward prediction using cells from Time 2 and Time 3 to infer the CGI distribution of Time 1. The predicted Time 1 N distribution is compared against the true distributions of Time 1-3. Right: Backward prediction using cells from Time 1 and Time 2 to infer the CGI distribution of Time 3, compared with the true distributions of Time 1-3. These results demonstrate that VAETracer accurately recovers generation-specific mutation accumulation patterns in both temporal directions.

(C) Comparison of predicted transcriptional latent states ($Z_{xt}$) with baseline methods on simulated data.

Left: Forward prediction of Time 1 using Time 2 and Time 3. UMAP visualization compares the predicted $Z_{xt}$ inferred by VAETracer with the true latent distributions of Time 1-3, alongside Time 1 latent representations inferred by LineageVAE. Right: Backward prediction of Time 3 using Time 1 and Time 2. Predicted $Z_{xt}$ from VAETracer is compared with the true Time 1-3 distributions and Time 3 latent representations inferred by WOT. VAETracer exhibits improved alignment with the true temporal structure in latent space.

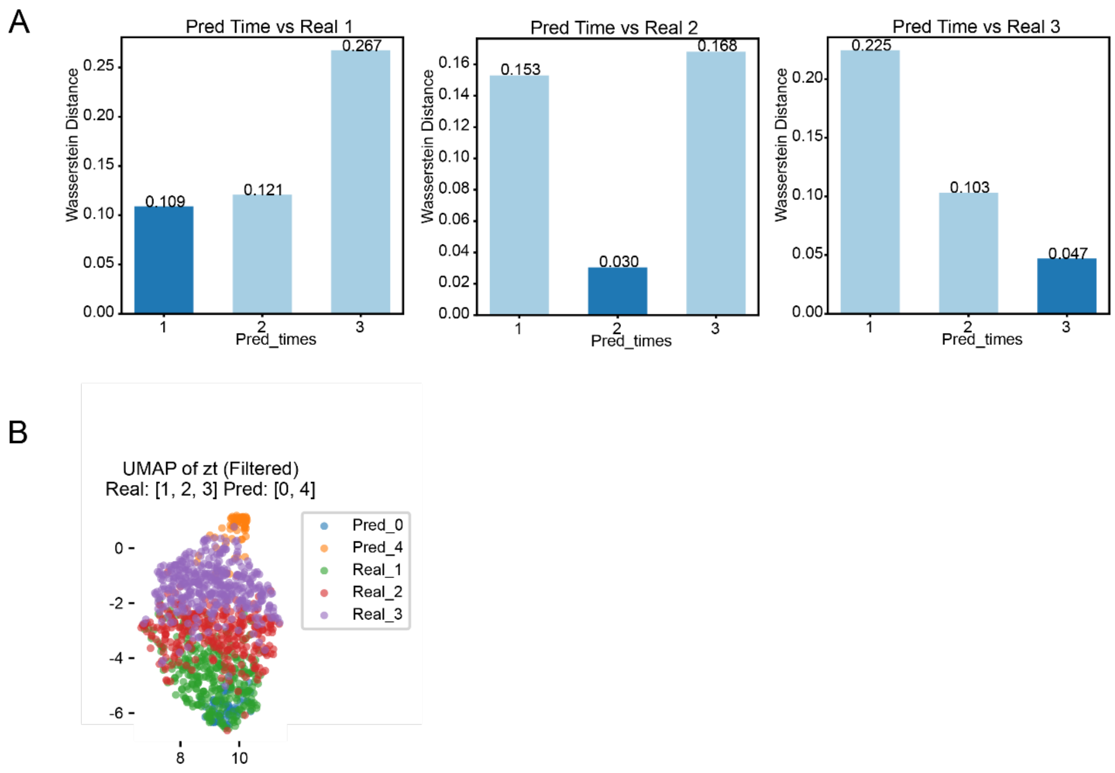


**Supplementary Figure 11: Validation on LUAS dataset.**

(A) Wasserstein distance between the predicted and observed CGI (N) distributions for LUAS cells under multiple inference settings: prediction of Time 1 using Time 2 and 3, prediction of Time 2 using Time 1 and 3, and prediction of Time 3 using Time 1 and 2. Lower distances indicate improved agreement between predicted and true generation distributions.

(B) UMAP visualization of predicted transcriptional latent representations ($Z_{xt}$) for unmeasured early (Time 0) and late (Time 4) tumor states inferred from Time 1-3 cells, together with the observed latent states of Time 1-3. The smooth spatial ordering supports coherent temporal extrapolation beyond measured time points.
